# Altered striatal dopamine dynamics and behavior in *Grin2a* mutant mice, a genetic mouse model of schizophrenia

**DOI:** 10.1101/2025.11.24.690326

**Authors:** Alyssa Hall, John Jacoby, Himanshu Gangal, Margery Chen, John Adeleye, Ethan Rochman, Nathaniel Shephard, Bryan Song, Zohreh Farsi, Sandeep Robert Datta, Morgan Sheng, Prabhat S. Kunwar

## Abstract

Schizophrenia (SCZ), a complex psychiatric disorder with a strong genetic basis, is thought to involve, at least in part, dopamine dysregulation in the striatum. Recent large-scale exome sequencing has identified multiple SCZ risk genes, including *GRIN2A* (encoding an NMDA (N-methyl-D-aspartate) receptor subunit) and *AKAP11* (encoding a protein kinase A binding protein), which is also a risk gene for bipolar disorder (BD). However, the mechanisms by which these genetic risk factors perturb dopamine circuits and cause psychotic symptoms remain poorly understood. We asked how behavior, dorsomedial striatal dopamine (DA) dynamics and the activity of D1-/D2-spiny projection neurons (SPNs) were altered in *Grin2a* and *Akap11* mutants. In the open field, *Grin2a* knockout mice display hyperlocomotion and an abnormal organization of naturalistic behaviors. They also displayed heightened behavioral responses to objects and auditory stimuli. *Grin2a* heterozygous animals mostly showed intermediate phenotypes, suggesting a dose-dependent effect of *Grin2a* loss. Further, *Grin2a* knockout mutants exhibited aberrant dopamine events, altered coupling of dopamine with locomotor features, and exaggerated stimuli-evoked dopamine responses. Notably, heterozygous animals also showed altered striatal SPN events in the open field reflecting dysregulated neural activity in both dSPNs and iSPNs. Compared to *Grin2a* mutants, *Akap11* mutants displayed opposite phenotypes, showing reduced locomotion with a shift from high- to low-velocity movement and distinct alterations in striatal neural activity. Together, *GRIN2A* (SCZ risk) and *AKAP11* (BD/SCZ risk) mutations induce different, often opposite, effects on striatal dopamine signaling and behavior, suggesting that these two risk genes act through distinct mechanisms.

## Introduction

Schizophrenia (SCZ) is a debilitating psychiatric disorder, part of a spectrum of severe mental illnesses that also includes bipolar disorder (BD), with a strong genetic basis that broadly impairs perception, motivation, cognition, and behavior^1–4^. Decades of neurobiological research have consistently implicated disturbances of dopaminergic signaling in the striatum as a core feature of the disease^5–7^. Positron emission tomography (PET) and other imaging studies, supported by pharmacological evidence, show increased amphetamine-evoked dopamine release, reflecting a hyperdopaminergic state, in the striatum in SCZ patients, which correlates with symptom severity and transition to psychosis^8,9^. Many SCZ symptom domains map onto striatal functions such as motivation, salience attribution, and motor control^7^, suggesting that dopaminergic dysregulation within striatal circuits may underlie certain clinical features. Despite this strong evidence, the mechanisms that give rise to dopaminergic circuit dysfunction and the resulting symptoms in SCZ remain poorly understood. Although SCZ and BD are clinically distinct, they share partially overlapping genetic risk, pointing to potential convergent pathways underlying these severe psychiatric illnesses^10,11^.

Recent large-scale exome sequencing studies, such as the Schizophrenia Exome Meta Analysis (SCHEMA) consortium, have identified rare, large-effect mutations^12^. Among these, *GRIN2A* is a SCZ-risk gene encoding a subunit of the NMDA receptor, which plays a critical role in glutamatergic neurotransmission^13,14^. Notably, *AKAP11* (encoding a protein kinase A (PKA) binding protein), a SCHEMA hit with a more modest association to SCZ, is a high-confidence risk gene for bipolar disorder^12,15,16^. Transcriptomic analysis from mutant mice shows that both *Grin2a* and *Akap11* mutations perturb gene expression and pathways in the spiny projection neurons (SPNs) of the striatum, a key region involved in dopamine signaling^13,17^. Consistent with those molecular disruptions, both *Grin2a* and *Akap11* showed heightened sensitivity to amphetamine in a test of amphetamine-induced hyperlocomotion^13,17^. Behavioral analyses have also identified multiple alterations, including markedly opposing locomotor phenotypes in *Grin2a* and *Akap11* mutants^13,14,17–21^. Together, these findings strongly suggest that rare SCZ- and BD/SCZ-associated mutations produce distinct and gene-specific disruptions in striatal dopamine circuitry and dopamine-modulated behavior.

Striatal dopamine circuits regulate motivation, motor action selection, and salience processing, behavioral domains consistently disrupted in SCZ^22–30^. Within the striatum, dopamine modulates striatal computations by modulating the activity of SPNs, which consist of direct (dSPNs) and indirect (iSPNs) output pathways expressing D1 and D2 receptors, respectively. These functions are anatomically segregated across subregions, including the dorsomedial (DMS), dorsolateral (DLS), ventral striatum, and tail of the striatum, each supporting specialized aspects of motivation, action, and salience processing^22,31,32^. Among these, the dorsomedial striatum (DMS), homologous to the associative striatum in humans, is particularly implicated in SCZ^7^ Compared to DLS, the DMS receives substantial frontal input and plays a central role in integrating cognitive, motivational, and sensorimotor signals to support flexible, goal-directed behavior and salience assignment^33–37^.

Few studies have directly examined how these SCZ- and BD-linked risk genes alter the striatal dopamine dynamics, SPN neural activity and associated behaviors. In this study, we integrated *in vivo* imaging of dopamine dynamics (via GRAB-DA) and SPN calcium activity (via GCaMP) in the DMS, with detailed, unsupervised machine-learning-based behavioral characterization of *Grin2a* and *Akap11* mutant mice. We measured both baseline locomotory activity and responses to external stimuli, domains known to strongly engage the dopaminergic system. Our results suggest specific effects of *Grin2a* loss-of-function on behavior, dopamine dynamics, and striatal SPN activity, which contrast with the behavioral and striatal SPN phenotypes of *Akap11* mutant animals.

## Results

### *Grin2a* knockout mice display hyperlocomotion and heightened behavioral responses to external stimuli

In previous reports, *Grin2a* mutant mice display several behavioral abnormalities, including hyperlocomotion^13,14,18–21^. We revisited and extended this behavioral characterization to include stimulus-evoked responses that engage salience processing mechanism, which is disrupted in schizophrenia patients^24,38–40^.

Consistent with prior findings, *Grin2a^-/-^* mice exhibited heightened velocities in a novel open field (**Fig. 1a-b**) compared to wild-type (WT) controls. However, animals across genotypes spent a similar amount of time in the inner zone of the open field (**Fig. 1c**), suggesting no change in anxiety-like (thigmotaxis) behavior. Upon subsequent introduction of a novel object at the center of the field (**Fig. 1a**), *Grin2a^-/-^*mutants exhibited marked increased interaction, as shown by multiple measures. They spent significantly more time in the center (**Fig. 1d**), made more frequent visits to the object (**Figure 1e**), and displayed a reduced latency to the first approach (**Fig. 1f**). While *Grin2a^+/-^* heterozygote animals did not differ significantly from WT in these measures, they showed intermediate values between wild type and knockout, consistent with a gene dose-dependent effect.

**Figure 1:**
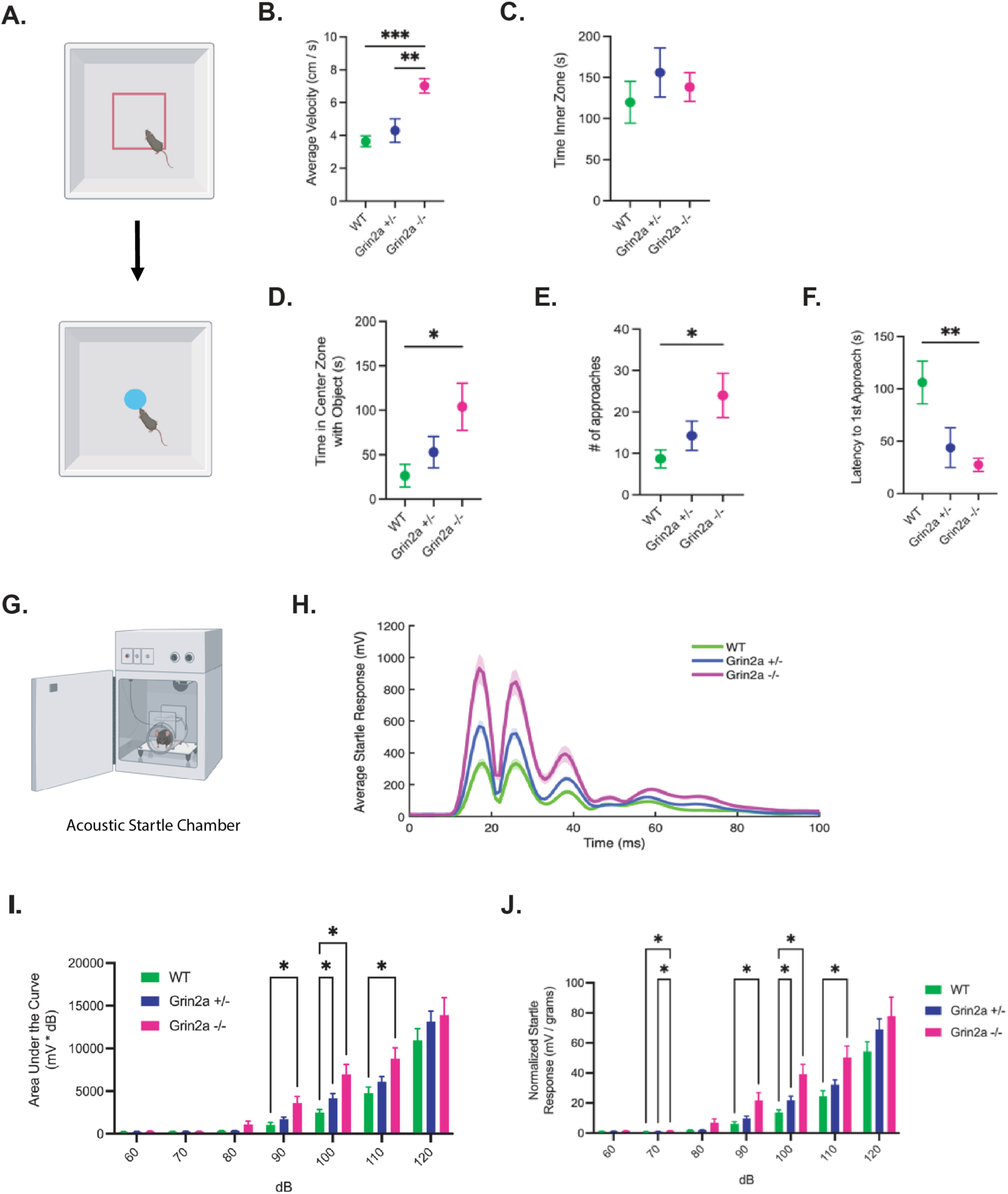
Grin2a knockout mice display hyperlocomotion and heightened behavioral responses to external stimuli **A.** Experimental design for open field and object interaction behavior recording. (n = 6 WT, 7 *Grin2a^+/-^*, 8 *Grin2a^-/-^* mice). **B.** Average velocity from 10-minute open field recording. **C.** Time in the inner zone of the open field. Inner zone drawn at 50% of the width of the open field. **D.** Time in the inner zone during object interaction. **E.** Number of approaches to the object during 10-minute recording. Approaches are defined entering a zone drawn around the object with a radius of 2 cm. **F.** Latency to first approach to the object. **B, C.** One-way ANOVA with Tukey’s post hoc test. *, p < 0.05; **, p < 0.01; ***, p < 0.001. **D, E, F.** Kruskal-Wallis Test with Dunn’s multiple comparisons test. *, p < 0.05. **G.** Schematic of acoustic startle chamber. **H.** Averaged waveform for acoustic startle response to 100 dB tone (WT: n = 80 trials, 10 mice; Grin2a^+/-^: n = 136 trials, 17 mice; Grin2a^-/-^: n = 72 trials, 9 mice). **I.** Area under the curve for initial startle response taken from 0-25ms following startle stimuli. **J.** Max startle response taken from averaged waveform from 0-25ms following startle stimuli and normalized to individual weights of mice. **I,J.** Repeated measures two-way ANOVA with Tukey’s post hoc test. *, p < 0.05. n = 10 WT, 17 Grin2a^+/-^, 9 Grin2a^-/-.^

To further characterize behavioral reactions to stimuli, we evaluated acoustic startle responses (**Fig. 1g-h**). *Grin2a^-/-^* animals showed an increase in startle responses across a range of sound intensities (60-120 dB) (**Fig. 1i, j**). The quantification of both area under the curve (AUC) of the motor force and its maximum peak within 25ms after the startle stimulus onset showed significant increases compared to WT animals (**Fig. 1i, j**). *Grin2a* heterozygotes also showed a marked increase in their startle response at 100 dB, with intermediate responses across tone levels, suggesting a gene-dose dependent impact (**Figure 1i-j**). The data also suggest a lower threshold for acoustic startle in *Grin2a-/-* mice, as they showed higher startle responses even to low decibel levels. However, unlike previous literature^19^, *Grin2a^-/-^ animals* did not show any differences in their prepulse inhibition. This suggests that while the magnitude of their startle response is heightened, their sensorimotor gating remains intact (**Extended Fig. 1e**).

*Grin2a* mutant animals were tested in spatial memory and sociability tasks. No significant differences in performance were observed in spontaneous alternation in the Y-maze (**Extended Fig. 1a**), but *Grin2a^-/-^* animals did display increased alternations and errors in this test (**Extended Fig. 1b, c**). Again, in the social interaction test, *Grin2a* mutant animals showed similar preferences for social interaction to those to WT animals (**Extended Fig. 1d**). In summary, *Grin2a* knockout mice exhibit hyperactivity in the open field, increased object interaction, and heightened responses to acoustic stimuli.

### *Grin2a* knockout mice display abnormal behavioral organization revealed by Keypoint MoSeq

We next examined whether loss of *Grin2a* alters the microstructure and temporal organization of spontaneous, solitary behavior in an open field. Beyond gross locomotory features such as velocity and acceleration, mouse behavior can be segmented into short, structured recurring segments/units of behavior known as “syllables”. These behavioral syllables can be classified into behavioral categories such as grooming, rearing, and pausing^26,41,42^. Further, the temporal organization of behavior can be quantified by examining the sequential structure of these syllables over time.

To further understand the behavioral organization of *Grin2a* mutants compared to WT controls, we recorded behavior for 60 minutes in empty Noldus Phenotyper boxes (**Fig. 2a**) (See Methods). The body pose estimation was done by a custom DeepLabCut model^43^, and the tracking pose data were analyzed by Keypoint MoSeq (KPMS)^42^.

**Figure 2:**
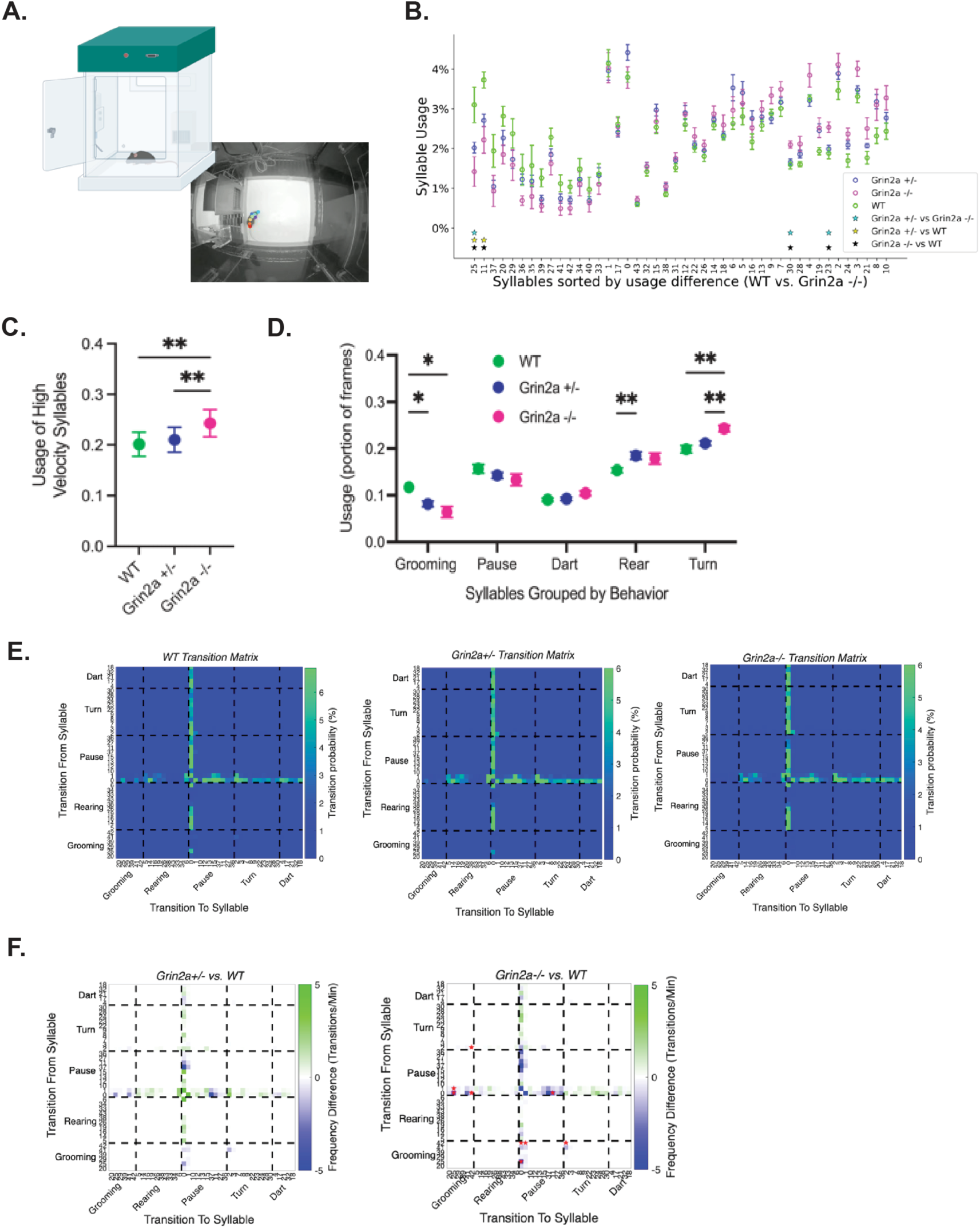
Grin2a knockout mice display abnormal behavioral organization revealed by Keypoint MoSeq **A.** Schematic of phenotyper recordings and example frame with DeepLabCut tracking. **B.** Syllable usage (proportion of frames) of WT and *Grin2a* mutant mice, sorted by usage difference between WT and Grin2a^-/-^ (n = 9 WT, 17 *Grin2a^+/-^*,9 *Grin2a^-/-^*). Stars on x-axis represent syllables with significant differences (FDR < 0.1) between genotypes. **C.** Syllable usage of syllables with velocities above the 75^th^ percentile of syllable velocity distribution (high velocity). One-way ANOVA with Tukey’s post hoc test. **, p < 0.01. **D.** Syllable usage grouped by annotated behavior groups. Two-way ANOVA with Tukey’s post hoc test. *, p < 0.05; **, p < 0.01. **E.** Heat map of transition matrices between syllables for WT, *Grin2a^+/-^* and *Grin2a^-/-^* animals, color bar represents scale of probability of transition. Syllables re-ordered according to behavioral groups. F. Heat map of subtraction of transition frequencies between WT vs. *Grin2a^+/-^*and WT vs. *Grin2a^-/-^.* Red stars indicate significance, FDR < 0.1.

*Grin2a* mutants displayed gene-dose-dependent differences in the usage across several syllables (**Fig. 2b**). The syllables with velocities in the top 25^th^ percentile were designated “high-velocity syllables”. Consistent with the previous report^13^. *Grin2a^-/-^* showed a significant upregulation in usage of high-velocity syllables (**Fig. 2c**). Manual annotation of syllables into behavioral categories revealed a reduction in usage of grooming-related syllables in heterozygous and homozygous animals in a dose dependent manner (**Fig. 2d**). *Grin2a^-/-^*animals also showed a notable increase in turning-associated syllables, and *Grin2a^+/-^* mutants displayed a significant increase in rearing syllables. These are consistent with enhanced exploratory drive (rearing, turning) and reduced self-maintenance (grooming) behavior in *Grin2a* mutants. To examine the sequential organization of syllables, we measured transitions between syllables. Transitions were organized by five behavioral groups: dart, turn, pause, rear, and grooming. In all animals, transitions were dramatically tied to pausing syllables, suggesting animals pause movement before switching to a different behavior (**Fig.2e**). *Grin2a* mutants displayed a reduction in transitions to and from pauses, consistent with more time spent moving (**Fig. 2f**).

Taken together, *Grin2a* knockout mice display altered use of behavioral “syllables”, with a marked shift from self-maintenance to exploratory syllables, along with abnormal transition frequencies between those syllables, while heterozygous animals showed intermediate phenotypes.

### *Grin2a* mutants exhibit altered dopamine dynamics during open field exploration

Given that *Grin2a mutants* display heightened locomotor activity and abnormal organization of syllables (**Figures 1 and 2**), we next examined whether dopamine (DA) dynamics in the dorsomedial striatum (DMS) are altered. Dopamine in dorsomedial striatum (DMS) plays critical roles in action selection, behavioral flexibility, and movement vigor^27,28,30,44,45^, and elevated dopamine in the striatum, particularly in the associative striatum (the human homolog of the rodent DMS), has been reported in the SCZ patients^8,9^. We therefore hypothesized that dopamine signaling in the DMS is dysregulated in *Grin2a* mutants. Specifically, we asked whether *Grin2a* mutations would alter baseline DA levels, the properties of spontaneous DA events (transient peaks above a certain threshold) or the coordination between DA dynamics and locomotor features such as acceleration, movement bouts, or pauses.

As an initial evaluation, we measured extracellular dopamine levels using microdialysis from the striatum before (baseline) and after Amphetamine (AMPH) administration in *Grin2a^+/-^* animals. Behaviorally, *Grin2a^+/-^* mice showed a trend towards an increased locomotor response to AMPH, consistent with the previous finding (**Extended Fig. 2a**)^13^. Following AMPH, all animals displayed the expected increase in dopamine level but with no difference in amplitude between WT, Grin2a^+/-^ and Grin2a^-/-^ mutants (**Extended Fig. 2b, e**). Baseline DA levels were also similar across genotypes (**Extended Fig. 2c-d**).

To measure DA dynamics with higher temporal resolution, we used fiber photometry to record in freely moving animals. We performed stereotaxic surgery on *Grin2a* and WT animals to inject the fluorescent DA reporter GRAB-DA using the syn promoter in AAV virus, in the DMS^46^ (**Fig. 3a**). GRAB-DA signals and locomotion were recorded in an open field for 60 minutes, simultaneously. As expected, *Grin2a^-/-^* animals remained hyperlocomotive during the recording, confirming the presence of this phenotype under this condition (**Extended Fig. 3a**). Next, we identified spontaneous DA events using an unbiased peak-detection algorithm and quantified each animal’s event frequency, event width, and event amplitude (**Fig. 3b**). *Grin2a^-/-^* animals showed a significantly increased event rate, about 4% higher than WT animals, and *Grin2a^+/-^*animals showed an intermediate phenotype (**Fig. 3c**). Width of GRAB-DA events was not significantly different across genotypes; though there was a reduction of event amplitudes in a gene dose-dependent manner, which did not reach significance (**Fig. 3c**).

**Figure 3:**
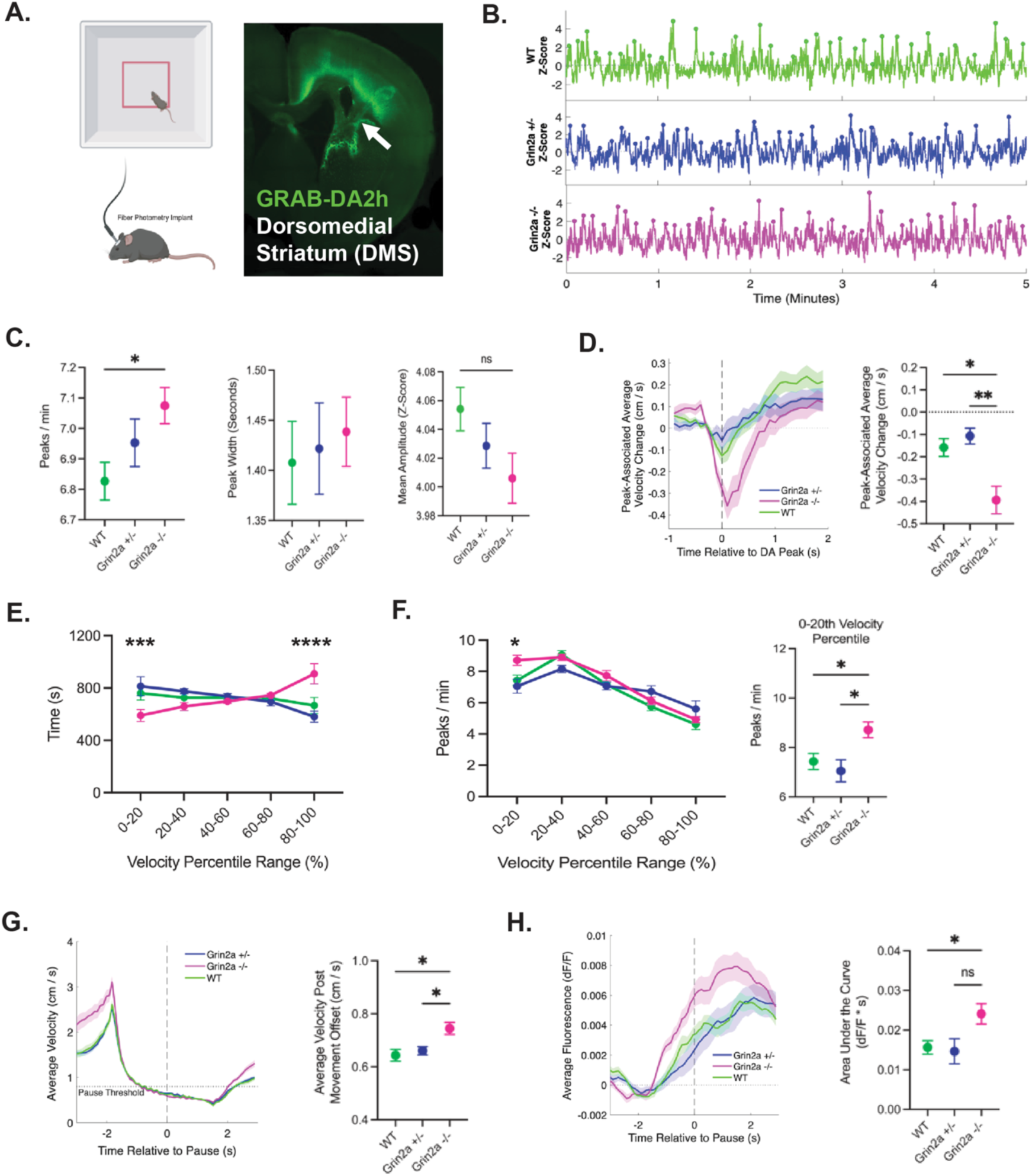
*Grin2a* mutants exhibit altered dopamine dynamics during open field exploration **A.** Schematic of experimental design for open field and dopamine fiber photometry recordings. Example histology of GRAB-DA2h injection in the dorsomedial striatum. **B.** Example z-scored GRAB-DA traces from WT (green), *Grin2a +/-* (blue), and *Grin2a -/-* (magenta) animals. Identified events (peaks) are marked with a circular point. **C.** Peak frequency, mean peak width, and mean peak amplitude of peaks identified from z-scored fiber photometry signal. N = 10 WT, 11 *Grin2a^+/-^*, 11 *Grin2a^-/-^*. **D.** Averaged velocity change associated with peak in WT and *Grin2a* mutants over time with SEM. Average velocity is normalized to 1s before peak onset to visualize change. Velocity change from 0-1s post peak is quantified between genotypes. **E.** Time spent in different velocity ranges. Velocity percentiles were calculated from the distribution of the velocities from all animals. To compare genotypes across bins a 2-way ANOVA was performed. A significant effect of velocity bin (p < 0.0001) was found. Using Tukey’s multiple comparison’s test, significant differences were found between WT and *Grin2a -/-* (p = 0.0135) and *Grin2a^+/-^* and *Grin2a^-/-^* (p = 0.0005) at velocities between the 0-20^th^ percentile. In the 80-100^th^ percentile, significant differences were also found between WT and *Grin2a^-/-^* (p = 0.0002) and *Grin2a^+/-^* and *Grin2a^-/-^* (p < 0.0001). **F.** Peak rates for different velocity ranges. Peak rates between genotypes shown for 0-20^th^ percentile of velocity. To compare genotypes across bins a 2-way ANOVA was performed. Significant effects of velocity bin (p < 0.0001), genotype (p = 0.0146), and the interaction of velocity bin and genotype (p = 0.0027) were found. Using Tukey’s multiple comparison’s test, significant differences between WT and *Grin2a^-/-^* (p = 0.0288) and *Grin2a^+/-^*and *Grin2a^-/-^* (p = 0.0179) were found at velocities between the 0-20^th^ percentile. A significant difference between WT and *Grin2a^+/-^* was found at velocities between the 20-40^th^ percentile (p = 0.0389). **G.** Averaged velocity for WT and *Grin2a* mutants over time during movement offset. Average velocity post-pause quantified between genotypes. **H.** Averaged dopamine signal over 6s during movement offset for WT and *Grin2a* mutants. Area under the curve is quantified from −1 to 3 seconds relative to beginning of movement offset. **C, D, F, G, H.** Comparisons between genotypes for a single measure were performed using Welch’s ANOVA test and Dunnet’s test for multiple comparisons. *, p < 0.05; **, p < 0.01.

To examine the relationship between GRAB-DA events and the animal’s movement in the open field, the average velocity was quantified at the time of DA events. In WT and *Grin2a^+/-^* animals, the timing of DA events corresponded with the maximum dip in a velocity reduction (**Fig. 3d**). *Grin2a^-/-^ animals* showed a dramatically larger velocity reduction at the time of dopamine events, suggesting that dopamine may have more influence over downstream motor behaviors in *Grin2a* mutants.

The analysis of DA event rates was aligned across common velocity bins defined by percentile ranges (0-20^th^, 20-40^th^, 40 – 60^th^, 60-80^th^, and 80-100^th^) that were calculated from the distribution of velocities from all animals. Interestingly, *Grin2a^-/-^*animals spent significantly less time in the lowest velocity bin and more time in the highest velocity, consistent with a hyperlocomotion phenotype (**Fig. 3e**). Across the genotypes, DA event rates were around seven events per minute, lasting over a second, suggesting a timescale slower than the sub-second duration of behavioral syllables identified by Keypoint MoSeq. Due to the difference in timescales and previous literature showing DMS dopamine does not correlate with syllable usage, it was not appropriate to directly align the dopamine signal with behavioral syllables^47^. In further analysis of the photometric signal, we saw DA event rates were higher in low velocity bins compared to high velocity bins (**Fig. 3f**), consistent with literature showing a negative correlation between DA and velocity^26,47–49^. *Grin2a^-/-^* mice showed a significantly increased rate of DA events in the lowest velocity bin, despite spending the least amount of time there (**Fig. 3f**). To further probe this relationship, we analyzed pauses (offsets) following the movement bouts to understand the average DA change associated with a reduction in movement. All animals showed increases in velocity after the pause, but *Grin2a^-/-^*animals showed higher velocities during this period (**Fig. 3g**). Notably, all animals showed an increase in average DA fluorescence corresponding with the reduction in movement, but this increase was significantly greater in *Grin2a^-/-^* animals (**Fig. 3h**). A complementary analysis involving movement onsets revealed that *Grin2a^-/-^* mice initiated movement at a higher velocity than controls (**Extended Fig. 3b**). The DA signals decreased during movement onset but *Grin2a*^-/-^ mice did not show a significant difference in their reduction compared to WT animals (**Extended Fig. 3c**).

Together, our results show that *Grin2a* knockout mice not only exhibit increased frequency of dopamine events but also display exaggerated dopamine signal and altered dopamine-movement coupling during pauses in movement. This altered dopamine signaling may contribute to the aspects of hyperlocomotion phenotype in *Grin2a* mutants.

### *Grin2a* knockout mice display exaggerated stimulus-evoked dopamine responses in the DMS

Environmental stimuli, such as novel objects, visual flashes or acoustic cues, evoke brief dopamine release in the DMS that signals novelty and salience^50–52^. Given that *Grin2a* mutants exhibit heightened behavioral reactivity to both object and acoustic stimuli, we next examined whether dopamine dynamics are altered during responses to these stimuli.

We first utilized GRAB-DA fiber photometry to record DA events during the presentation of simple, non-contingent sensory cues. Mice received 20 brief light flashes and 20 brief tones (**Fig. 4a, e**), delivered in four blocks of five stimuli, to assess both initial response magnitude and habituation across repeated presentations. In WT animals, DA responses were phasic and sharply increased around ∼0.5 seconds following the onset of the light cue and their amplitude declined across successive presentations (**Fig. 4b, c**)., *Grin2a^-/-^* showed significantly larger and more prolonged responses than WT, especially during the first time block (**Fig. 4c**). Quantification of the average area under the curve (AUC, 0-4 seconds post stimuli onset) across the five stimuli per block confirmed *Grin2a^-/-^* had a higher novel cue-evoked DA response for the first trial block but habituated similarly to WT animals across subsequent trial blocks (**Fig. 4d**). For auditory stimuli, a similar pattern was observed in that *Grin2a^-/-^* animals showed an elevated response to the first block of stimuli (**Fig. 4f-h**). However, the response to auditory cues was much lower and variable across animals. Notably, the heterozygous animals showed intermediate phenotypes in the first stimulus block.

**Figure 4:**
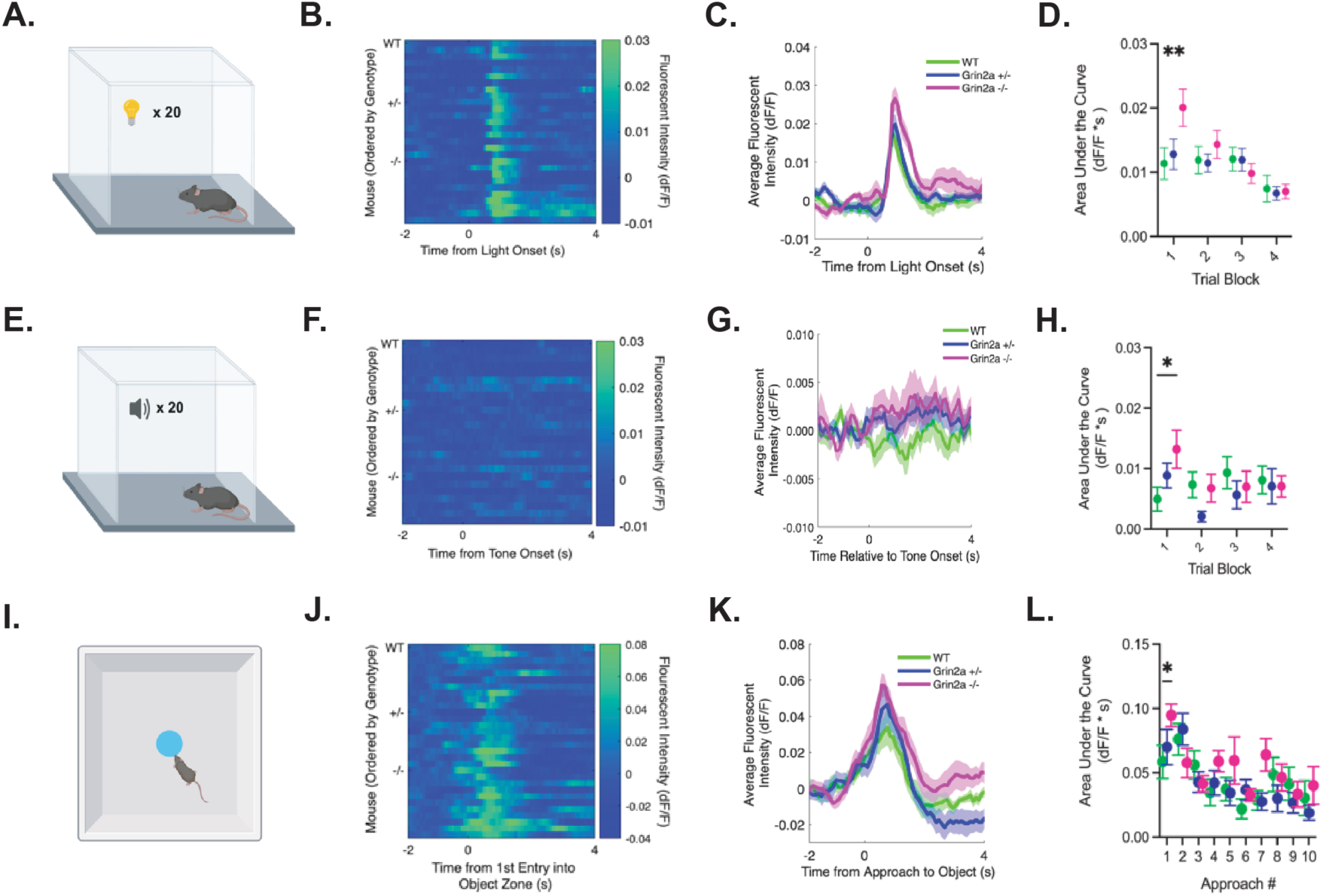
*Grin2a* knockout mice display exaggerated stimulus-evoked dopamine responses in the DMS **A.** Schematic of experimental setup for light presentations. **B.** Heat map of averaged dopamine response from first block of light presentations (five trials per block) from each subject (ordered by genotype). N = 9 WT, 10 *Grin2a^+/-^*,10 *Grin2a^-/-^*. **C.** Averaged dopamine responses by genotype from first block of light presentations. Normalized to baseline period of −2 to −1s before light onset. **D.** Area under the curve calculated from 0 to 4s post light onset compared between genotypes for each trial block of light presentations. **E.** Schematic of experimental setup for tone presentations. **F.** Heat map of averaged dopamine response from first block of tone presentations (five trials per block) from each subject (ordered by genotype). N = 9 WT, 10 *Grin2a^+/-^*,10 *Grin2a^-/-^*. **G.** Averaged dopamine responses by genotype from first block of tone presentations. Normalized to baseline period of −2 to −1s before tone onset. **H.** Area under the curve calculated from 0 to 4s post light onset compared between genotypes for each trial block of light presentations. **I.** Schematic of experimental setup for object interaction. **J.** Heat map of averaged dopamine response from first approach to object (five trials per block) from each subject (ordered by genotype). N = 10 WT, 10 *Grin2a^+/-^*,10 *Grin2a^-/-^*. **K.** Averaged dopamine responses by genotype from first approach to object. Normalized to baseline period of −2 to −1s before entry into object zone. **L.** Area under the curve calculated from 0 to 2s post light onset compared between genotypes for first 10 approaches to novel object. **D, H,L.** Repeated measures 2-way ANOVA with Tukey’s post hoc test. *, p < 0.05; **, p < 0.01.

To examine DA responses to a more naturalistic salient stimulus, we next recorded DA dynamics during interaction with a novel object (**Fig. 4i**) over a 25-minute session. During the initial approach to the object, *Grin2a^-/-^* showed significantly higher dopamine responses compared to WT animals (**Fig. 4j, k**). Quantification of area under the curve (AUC) (0-2s post entry to object zone) across the first 10 approaches confirmed a significant increase to the first approach, but not to subsequent approaches (**Fig. 4l**). Behaviorally, *Grin2a^-/-^* animals showed a significant increase in approach frequency during the first 10 minutes, but the effect was not significant in the rest of the recording, aligning with the DA measurement data (**Extended Fig. 4a**).

Taken together*, Grin2a* knockout mice display increased stimulus-evoked dopamine activity in the DMS in response to novel stimuli, while heterozygotes show intermediate impact. These findings identify changes in DMS dopamine dynamics that parallel the increased behavioral reactivity to stimuli observed in *Grin2a* mutants.

### Altered neural activity in striatal direct and indirect pathway neurons in *Grin2a* heterozygous mice

After demonstrating that *Grin2a* mutations alter DMS dopamine dynamics and behavior, we next examined whether these changes extend to downstream circuit activity. Dopamine signaling to D1- and D2-SPNs and their coordinated activity is essential for regulating movement and salience-related behaviors, but dopamine acts on these cell types in opposing ways^26,44,53,54^. Specifically, elevated dopamine levels increase activity of D1-SPNs but decreases the activity of D2-SPNS^55,56^. We used *Grin2a* heterozygous mice, which show intermediate behavioral as well as dopamine phenotypes, to unravel the higher resolution cell type-specific effects in SPNs, which possess either D1 (dSPNs) or D2 receptors (iSPNs). Given that human SCZ patients with a *GRIN2A* mutation only carry one copy of the genetic mutation, this approach more directly reflects the human genetic condition^12^.

To measure cell-type specific differences in the DMS, we employed calcium imaging using fiber photometry (**Fig. 5a**). A Cre-dependent AAV GCaMP virus was injected into WT and *Grin2a^+/-^ mice* crossed with D1-Cre and A2A-Cre mice to record from dSPNs and iSPNs, respectively. In addition, for the population-level photometric measurements, a non-specific calcium reporter AAV virus (CaMKII-GCaMP) was injected in WT and *Grin2a^+/-^*mice.

**Figure 5:**
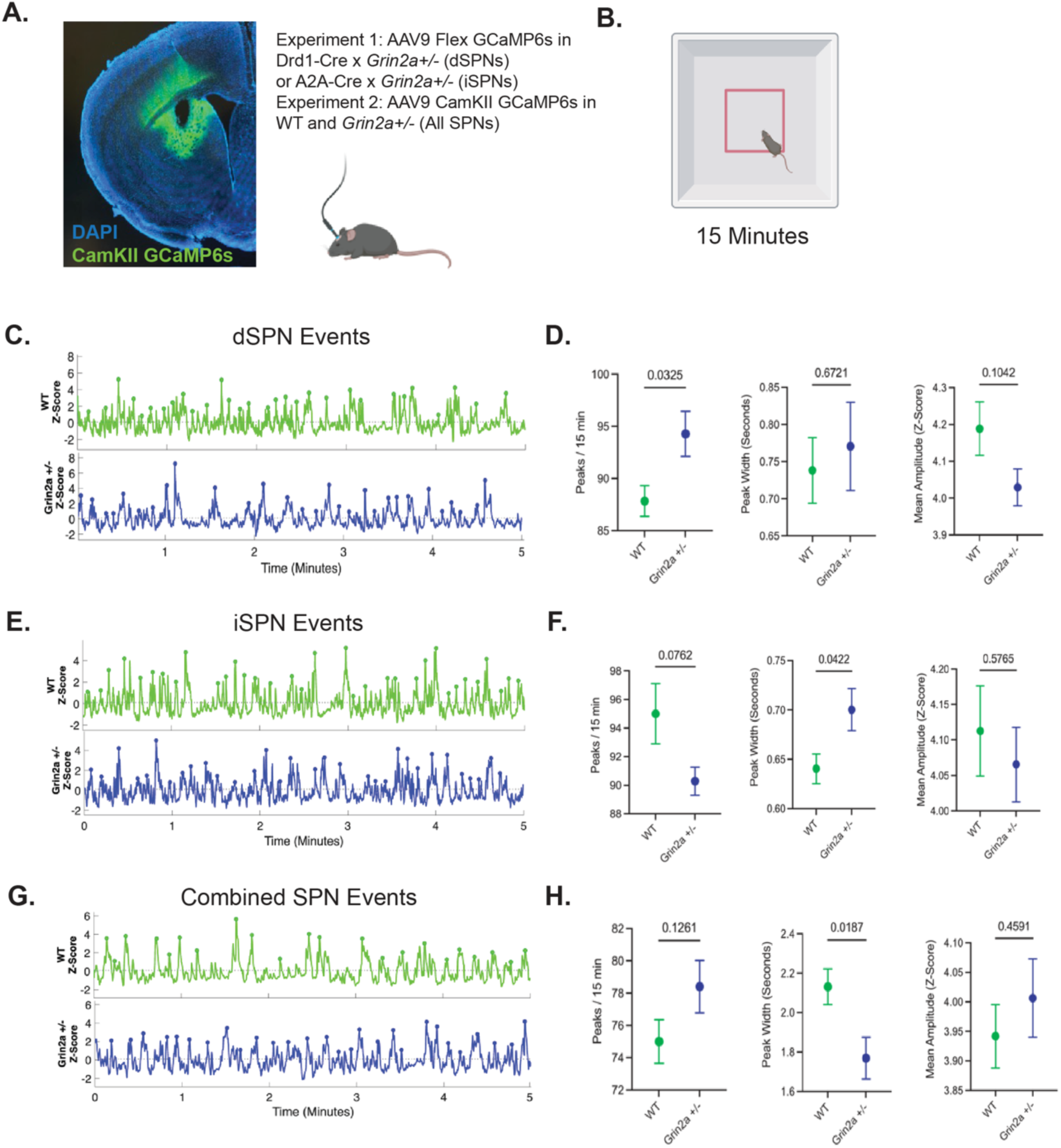
Altered neural activity in striatal direct and indirect pathway neurons in *Grin2a* heterozygous mice **A.** Example histology of GCaMP injection in the dorsomedial striatum. Outline of injections in different strains. **B.** Schematic of open field recording. **C.** Example traces from D1-Cre (WT and *Grin2a^+/-^*) animals, measuring dSPN activity. Identified events (peaks) are marked with a circular point. **D.** dSPN peak characteristics of WT and *Grin2a^+/-^* animals (frequency, mean peak width, amplitude). N = 6 WT and 7 *Grin2a^+/-^*. **E.** Example traces from A2A-Cre (WT and *Grin2a^+/-^*) animals, measuring iSPN activity. Identified events (peaks) are marked with a circular point. **F.** iSPN peak characteristics of WT and *Grin2a^+/-^* animals (frequency, mean peak width, amplitude). N = 7 WT and 7 *Grin2a^+/-^*. **G.** Example traces from WT and *Grin2a^+/-^* animals, measuring total calcium activity in the DMS. Identified events (peaks) are marked with a circular point. **H.** Bulk calcium peak characteristics of WT and *Grin2a^+/-^* animals (frequency, mean peak width, amplitude). N = 9 WT and 10 *Grin2a^+/-^*. **D., F, H.** Comparisons were made using Welch’s T test. *, p < 0.05.

We first quantified calcium events reflecting neural activity during 15-minute baseline recording sessions in an open field (**Fig. 5b-h**). *Grin2a^+/-^* mice exhibited a 5% significant increase in dSPN calcium event frequency compared to WT mice (**Fig. 5d**) and nonsignificant changes in mean event width and event amplitude, although there was a trending decrease for event amplitude. Conversely, *Grin2a^+/-^*mice showed a modest, non-significant decrease (∼5%) in iSPN event frequency (**Fig. 5f**) but a significant increase in mean event width when compared with WT controls (**Fig. 5f**). In CaMKII-GCaMP recordings, the photometric recordings revealed a decrease in event width and a non-significant trend towards increased event frequency in *Grin2a^+/-^*(**Fig. 5h**). None of the event-related analyses showed differences in mean event amplitude (**Fig.5d,f,h**).

Next, we examined how SPN activity is related to locomotion in the open field (**Extended Fig. 5a**) by aligning calcium activity to velocity, movement onset, and pause onset in *Grin2a^+/-^* mutants. Both dSPNs and iSPNs showed similar activity patterns across velocity bins (**Extended Fig. 5b**) and during movement onset (**Extended Fig. 5c**) and offset (**Extended Fig. 5d**). *Grin2a^+/-^* animals showed comparable activity profiles across velocity bins (**Extended Fig. 5e, h**), at movement onset (**Extended Fig. 5f, i**), but exhibited a nonsignificant decrease in the signal change following the pause at movement offset in both dSPNs and iSPNs (**Extended Fig. 5g, j**). Similarly, the photometric recordings from all SPNs showed similar a similar increase in event rates at different velocities and an increase in activity at movement onset (**Extended Fig. 5k-l**), but there were no differences in these measures in *Grin2a^+/-^* animals. During pauses (movement offsets), *Grin2a^+/-^* animals did show a significant reduction in total SPN activity (**Extended Fig. 5m**). In alignment of velocity to the time of SPN events, the change in velocity was variable across dSPNs and iSPNs, but combined SPN activity showed a net decrease in velocity following event onset. Although there were no significant differences observed between genotypes in the velocity change associated with the time of event onset (**Extended Fig. 5n**).

These results suggest that in *Grin2a^+/-^* mutants, there is a modest shift of SPN calcium activity (GCaMP signal) to favor a higher D1/D2 activity ratio, driven by increased dSPN activity relative to iSPN activity. Notably, the SPN activity changes observed in *Grin2a^+/-^* mice align with what we would expect given their slightly increased dopamine response (**Fig. 3c**), potentially suggesting a coordinated disruption of both dopaminergic input and downstream striatal output pathways.

### *Akap11* mutant animals display opposite alterations in locomotion and striatal neural activity

As a comparison against *Grin2a* mutants, we investigated mice lacking *AKAP11*, a strong bipolar disorder and modest SCZ risk gene that encodes a Protein Kinase A (PKA) anchoring protein^12,15–17,57^. Importantly, *Akap11* mutant mice exhibit hypolocomotion^17,18,57^, which is opposite to the hyperlocomotion observed in *Grin2a* mutants. We hypothesized that *Akap11* mutations perturb striatal dopamine signaling in the opposite direction to *Grin2a* mutants, leading to reduced locomotor drive and altered striatal neural activity.

We first analyzed locomotor behaviors in *Akap11* mutants in an open field using conventional methods and the Keypoint-MoSeq pipeline (**Fig. 6a**). *Akap11^-/-^* mice displayed a significant reduction in total distance traveled over a 60-minute open field recording, confirming a hypolocomotion phenotype reported (**Fig. 6b**). *Akap11^+/-^* generally showed an intermediate phenotype. Keypoint-MoSeq analysis revealed altered syllable usage patterns (**Fig. 6c**). When grouped by annotated behavior, *Akap11^-/-^* showed significantly reduced use of rearing-related syllables and an increase in pausing-related syllables (**Fig. 6d**). A reduction in rearing suggests less exploratory behavior in *Akap11* mutants and is opposite to the increase in rearing we observed in Grin2a heterozygous animals (**Fig. 2d**). Furthermore, transition analysis did not reveal significant differences between WT and *Akap11* animals, but *Akap11* knockout animals did show a trending increase in pause-related transitions (**Extended Fig. 6a, b)**).

**Figure 6:**
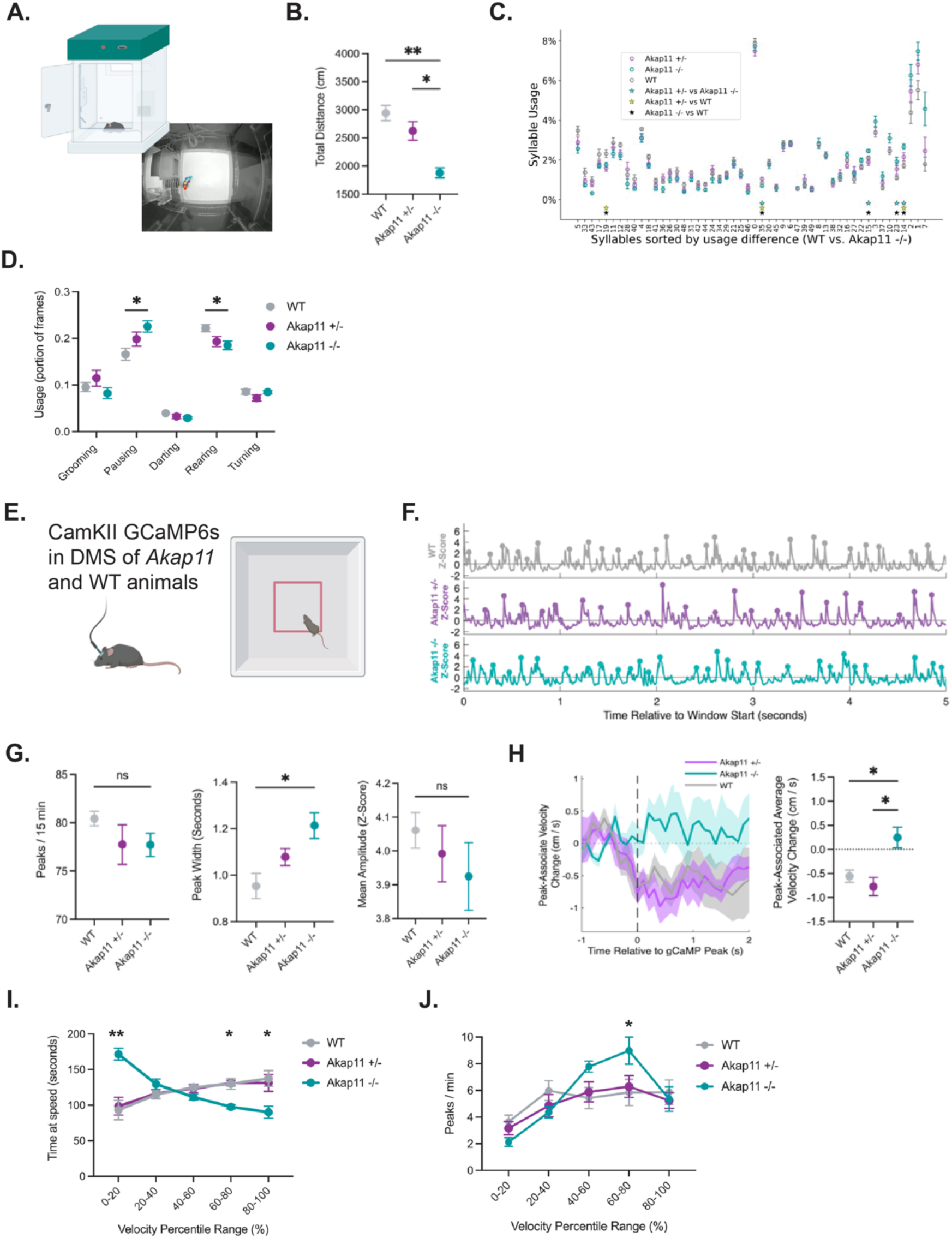
*Akap11* mutant animals display opposite patterns in locomotion and striatal neural activity **A.** Schematic of phenotyper recordings and example frame with DeepLabCut tracking. **B.** Total distance travelled measured over a 60-minute open field recording between WT and *Akap11* mutants. N = 8 WT, 9 *Akap11^+/-^*, 7 *Akap11^/-^*. **C.** Syllable usage (proportion of frames) of WT and *Akap11* mutant mice, sorted by usage difference between WT and *Akap11 -/-.* (n = 9 WT, 17 *Akap11^+/-^*, *9 Akap11^-/-^*). Stars on x-axis represent syllables with significant differences (FDR < 0.1) between genotypes. **D.** Syllable usage grouped by annotated behavior groups. Two-way ANOVA with Tukey’s post hoc test. *, p < 0.05. **E.** Schematic of fiber photometry and open field recordings. **F.** Example traces from WT, *Akap11^+/-^* and *Akap11^-/-^*. animals, measuring total calcium activity in the DMS. Identified events (peaks) are marked with a circular point. **G.** Bulk calcium peak characteristics of WT, *Akap11^+/-^* and *Akap11^-/-^* (frequency, mean peak width, amplitude). N = 7 WT and 8 *Akap11^+/-^* and 7 *Akap11^-/-^*. **H.** Averaged velocity change associated with peak in WT and *Akap11* mutants over time with SEM. Average velocity is normalized to 1s before peak onset to visualize change. Velocity change from 0-1s post peak is quantified between genotypes. Comparisons made using Welch’s t test. *, p < 0.05. **I.** Time spent in different velocity ranges. Velocity percentiles were calculated from the distribution of the velocities from all animals. To compare genotypes across bins a 2-way ANOVA was performed. A significant effect of the interaction of velocity bin and genotype (p < 0.0001) was found. Using Tukey’s multiple comparison’s test, significant differences were found between WT and *Akap11^-/-^* in the ranges of the 0-20^th^ percentile, 60-80^th^ percentile, and 80-100^th^ percentile. **J.** Peak rates for different velocity ranges. Peak rates between genotypes shown for 0-20^th^ percentile of velocity. To compare genotypes across bins a 2-way ANOVA was performed. A significant effect of the velocity bin (p < 0.0001) was found but a significant effect of the interaction of velocity and genotype was not present (p = 0.0517). Using Tukey’s multiple comparison’s test, significant differences between WT and *Akap11^-/-^*(p = 0.0108) and *Akap11^+/-^*. and *Akap11^-/-^* (p = 0.0271) were found at velocities between the 60-80^th^ percentile.

To explore the underlying physiologic changes, we performed bulk calcium imaging (CaMKII-GCaMP) in the DMS during open field behavior (**Fig. 6f, g**). *Akap11* mutants exhibited a trend towards decreased calcium event frequency and a significant increase in event width compared to WT animals (**Fig. 6g**), indicating wider and trending towards less frequent activity in the DMS, opposite to the increased frequency and reduced width observed in *Grin2a* mutants (**Fig. 3c**).

We next analyzed the coupling between DMS calcium events and locomotor state in *Akap11* and WT mice. When aligning velocity to calcium event onsets, WT and *Akap11*^+/-^ showed a decrease in average velocity at the time of calcium events, but *Akap11^-/-^* animals did not (**Fig. 6i**). Velocity-bin analysis revealed that, as opposed to Grin2a^-/-^ (**Fig. 3e**) *Akap11^-/-^* mice spend significantly less time in the highest velocity bins (60-100^th^ percentile) and more time in the lowest velocity bin (0-20^th^ percentile) compared to WT mice, confirming a shift towards slower movement (**Fig. 6i**). Notably, *Akap11*^-/-^ animals displayed a significantly increased calcium event rate during faster movement bouts (60-80^th^ percentile velocity bin) compared to both *Akap11^+/-^* and WT animals (**Fig. 6j**).

Movement-bout analysis in *Akap11* mutants versus WT further showed nonsignificant trends of a lower maximum velocity achieved during movement onset (**Extended Fig. 7a**) and reduced average velocity at movement offset (**Extended Fig. 7b**). In *Akap11* mutants, calcium dynamics showed a trend toward increased activity at movement onset (**Extended Fig. 7c**) and a decrease in activity during movement offset (**Extended Fig. 7d**), compared to WT animals.

Taken together, *Akap11* mutations lead to hypolocomotion, with a shift to lower velocity, reduced exploratory syllable usage, slower and broader striatal calcium events, and an aberrant relationship of striatal calcium activity to velocity. These findings contrast with those related to the *Grin2a* mutants and suggest that loss of *AKAP11* (linked to bipolar and SCZ) or *GRIN2A* (linked to SCZ) result in different, often opposite, changes in the striatal activity and behavior.

## Discussion

### *GRIN2A* (SCZ risk): dopamine signaling and behavior

*Grin2a* knockout mice exhibited both increased dopamine events and exaggerated novel cue-evoked responses, which is consistent with human evidence of a hyperdopaminergic state and aberrant salience in SCZ patients. In parallel, behavior measures that often engage dopamine signaling, including locomotion and responses to novel stimuli, were similarly heightened. One can speculate a potential causal relationship between elevated dopamine activity and behavioral reactivity, which can be tested by circuit-level interventions. Notably, *Grin2a* heterozygous mice showed intermediate alterations across dopamine dynamics, behavioral measures, and D1/D2 SPN neural activity, including a modest shift towards D1-SPN activation. These heterozygous phenotypes were often mild or below the threshold for statistical significance, reflecting subtle abnormalities rather than overt deficits. This finding aligns with human genetics, in which SCHEMA-identified variants are heterozygous loss-of-function mutations^12^. It suggests that the partial loss of NMDAR function could reflect a “risk-state” in which additional genetic or environmental factors such as developmental stress or drug exposure are required to unmask the latent state to clinically relevant circuits and behavioral abnormalities^58^.

### *AKAP11* (BD/SCZ risk) and contrast to *GRIN2A* (SCZ risk)

*Akap11* mutant mice exhibit reduced locomotion, a shift toward low-velocity movement, and diminished SPN calcium activity. These patterns stand in clear contrast to the hyperlocomotive and elevated dopaminergic phenotypes in *Grin2a* mutants. This directionality of the *Akap11* phenotype could reflect its stronger genetic association with bipolar disorder relative to schizophrenia, raising the possibility that *AKAP11* disturbance could model hypoactive, hypodopaminergic circuit states relevant to mood dysregulation of bipolar disorder (BD)^12,15^. Although substantial further work will be needed to test this hypothesis directly, our findings suggest that a balance of striatal dopaminergic signaling may represent a shared vulnerability that is common between SCZ and BD, with respective genes biasing dopamine signaling in opposite directions.

### Limitations

Several limitations remain in our study. First, a direct and similar experimental comparison of dopamine and D1-/D2-SPN signals between *Akap11* and *Grin2a* mutants will be necessary to clearly address their shared and divergent circuit impacts. Future work extending these analyses to additional SCZ- and BD-risk genes will be vital. Our recordings were restricted to the dorsomedial striatum and based on bulk fiber photometry, which cannot resolve temporal and single cell activity or capture dynamics across other striatal nodes such as DLS, ventral striatum or midbrain dopamine neurons^26,44,53,59^. Further, the heterozygous phenotypes we studied were modest and remain incompletely characterized. Stress exposure or other demanding cognitive or motivational tasks may be needed to unravel latent circuit or behavioral disruptions. Importantly, future research must explicitly include female subjects to determine whether the *Grin2a* and *Akap11* mechanisms generalize or display sex-specific vulnerability.

## Acknowledgements

We thank Tyler Caron, Matthew Beck, Nathan Chan, James Ingersoll, and Leiby Vargas for help with mouse colony management and logistics for behavioral and fiber photometry experiments; Members of the Sheng lab for valuable feedback on the project; John Adeleye for assistance with perfusions; Members of the Granger lab, Fu lab, and Pan lab, specifically Kate Stalnaker and Sean Moran, for sharing of behavioral space and protocols; Jones Parker for guidance on in vivo recordings in dSPNs and iSPNs; Katie Heslip and Natasha Kulviwat for assistance in histology; Yaqi Li for assistance in labeling frames for the initial DeepLabCut model; Dana Levy, Ann Kennedy, Aditya Nair for guidance of analysis of the fiber photometry data.

## Author Contributions

M. S. and P.S.K. conceived of the project and supervised all aspects of the work. A.H. designed, conducted, and analyzed all experiments. J.J. assisted with the Keypoint MoSeq data analysis under the supervision of S.R.D., who also advised on the project. A.H., H.G., and N.S. set up the fiber photometry recording system under guidance of P.S.K. M.C. and J.A. contributed to the histological analyses. H.G. assisted in development of scripts for event-analysis of fiber photometry experiments and testing stimulus-evoked DA responses. E.R. assisted in the analysis of behavior experiments. B.S. and Z.F. contributed reagents and assisted with experimental design. P.S.K. and A.H. wrote the manuscript with assistance from M.S. and S.R.D., and with input from all authors.

## Methods

### Animals

All experiments were approved by the Broad Institute IACUC (Institutional Animal Care and Use Committee) and conducted in accordance with the NIH Guide for the Care and Use of Laboratory Animals. *Grin2a^+/-^, Grin2a^-/-^,* and wild-type littermates were generated and crossed as previously described^13^. Drd1-Cre (B6.FVB(Cg)-Tg(Drd1a-cre)262Gsat/Mmcd) and A2A-Cre (B6.FVB(Cg)-Tg(Adora2a-cre)KG139Gsat/Mmcd ET5568) mice were were obtained from the UC Davis Mouse Biology Program (MMRC) and were crossed against wild-type C57/BL6J mice (Jackson Laboratory, #000664). The Drd1-Cre and A2A-Cre mice were then further crossed with *Grin2a^+/-^* mice to generate *Drd1-Cre; Grin2a^+/-^, Drd1-Cre; Grin2a +/+, A2A-Cre; Grin2a^+/-^,* and *A2A-Cre; Grin2a^+/+^* animals. *Akap11^+/-^, Akap11^-/-^*, and wild-type littermates were generated and crossed as previously recorded (Song et al., 2025). For all experiments, male adult mutant mice and wild-type littermates, between 4-6 months, were used.

### Behavioral Tests

#### Open Field Test

For open field tests, mice were tested in a plexiglass box (Noldus, 40×40×30 cm) and behavior was recorded overhead by a camera (Basler Color GigE Camera, Noldus). Animals were habituated to the behavioral room for at least thirty minutes prior to the recording. Recording time was kept consistent for mice in each experimental cohort and between mice, the arena was cleaned with a Clidox-S solution. Metrics of velocity and distance travelled were determined from automated tracking with Ethovision. A square center zone was drawn in the center of the arena with a diameter of 50% of the width of the open field arena. Time spent in the center zone was calculated and compared across genotypes.

#### Object Interaction

Measurement of object interaction was performed in the open field boxes (Noldus, 40×40×40 cm) and behavior was recorded overhead by a camera. Prior to object introduction, animals were recorded in the open field for 10 minutes to allow for habituation. The object used was a small, clear glass (Diameter = 4 cm, Height = 5 cm; Ikea) and was placed upside down so that animals could not climb inside the object. During the recording, the object was placed in the center of the arena and between animals, was cleaned with Clidox-S solution. Ethovision was used to track behavior and approaches to the object were defined by entering a zone with a radius of 2cm away from the perimeter of the object. Frequency of approaches was defined as the # of approaches divided by the duration of the recording.

#### Acoustic Startle and Prepulse Inhibition

Acoustic startle response (ASR) measurements were performed using the SR-LAB™ Startle Response System (San Diego Instruments). Individual mice were tested in five startle apparatuses consisting of a plexiglass, ventilated, cylindrical holding tube (5” L x 1.5” ID) within a sound attenuated chamber (15” W x 14” D x 18” H). One day before testing, mice were placed into the holding tube inside the recording chamber for 15 minutes to allow for habituation. On the day of the experiment, a 60 dB background tone was played throughout the recording. Prior to recordings of startle responses, mice had a five-minute acclimation period with only the background tone. Following acclimation, mice were exposed to 56 trials of varying decibel levels (60, 70, 80, 90, 100, 110, 120 dB; 40ms, 8 trials each) with pseudorandom inter-trial intervals of 10-30s. Startle responses were measured with a piezoelectric accelerometer underneath the holding tube in mV for 25 ms following the startle stimuli and then normalized to the corresponding animal’s weight.

PPI evaluation occurred one week after acoustic startle measurements and included a five-minute acclimation period followed by five priming acoustic stimuli (110 dB, 40ms). After the priming stimuli, thirty trial blocks were presented in a pseudo-random order with either no stimulus or a pre-pulse tone (65, 70, 75, 80, 85 dB; 20 ms). The prepulse occurred 100 ms before the startle stimulus (110 dB, 40 ms). Following the prepulse trials, another block of five startle stimuli was delivered (110 dB, 40 ms). The average response to 110 dB was calculate from the first block and last block of startle presentations, without prepulse stimuli, and were normalized to the corresponding animal’s weight. Prepulse inhibition was calculated by the following formula: [average normalized startle response (110 dB) – normalized startle response with prepulse] / [average normalized startle response (110 dB)] x 100%.

#### Spontaneous Alternation in the Y-maze

Testing for spontaneous alternations was performed in a symmetrical maze with three identical arms (35×5×20 cm, Noldus). To visually distinguish arms, a different shape was drawn on a flat paper taped at the top of each arm (circle, square, or triangle). At the start of the test, each mouse is placed at the end of one arm, furthest from the center. Animals were allowed to explore the maze for 20 minutes and the initial arm was alternated across tests. Correct alternations were scored when the mouse visits each arm sequentially without revisiting any of the arms. To calculate the percentage of correct or “spontaneous alternations”, we used Ethovision tracking and calculated the number of correct alternations divided by the maximum number of possible correct alternations and multiplied by 100%.

#### Three Chamber Social Interaction Test

The social interaction apparatus consisted of three equally sized chambers: one left, one central, and one on the right, in a plexiglass arena (40.5×60×22 cm, Noldus). Mice were acclimated to the test room for thirty minutes prior to testing. After acclimation, the mouse was placed in the center chamber and allowed to freely explore the empty 3 chamber box for 5 minutes. After 5 minutes, the mice were removed from the box or confined to the center chamber. In trial 1, an acrylic cup with wires (10cm diameter, 20cm height) containing a novel adult sex matched mouse was placed in one of the other two chambers. A second empty acrylic cup with a wire cup was placed in the opposite chamber. The dividers were then lifted so that the mouse could freely move through all three chambers. The mouse was then allowed to freely explore for 10 minutes. After these 10 minutes, in trial 2, a second novel adult sex matched mouse was placed in the empty acrylic cup. The mouse was allowed to freely explore for another 10 minutes. The chamber the novel mouse is placed in was alternated between the left and right chambers to account for a chamber preference in the test mice. The social interaction arena was cleaned with Clidox-S solution in between animals The amount of time spent exploring the novel mouse and the time spent exploring the empty cup was tracked using Ethovision. The discrimination index for each trial (social vs. empty and novel vs. familiar mouse) was calculated by the following formula [time spent with chamber of interest / time spent with other chamber].

### DeepLabCut and Keypoint MoSeq Pipeline

Keypoint MoSeq is an unsupervised machine learning tool for identifying behavioral modules (“syllables”) from key point tracking data^42^. For pose estimation and key point labeling, we trained DeepLabCut models^43^ on top-down videos recorded at 25 fps with ResNet-50. The *Grin2a* model was trained on 400 manually annotated frames spread across 20 videos and the *Akap11* model was trained on 300 manually annotated frames spread across 15 videos. Training frames were split into 95% used for training and 5% used for testing. Each model was trained over 100,000 iterations. The key points included were the right ear, left ear, and six points along the dorsomedial axis of the mouse’s body. For fiber photometry recordings, we trained a separate deeplabcut model (300 training frames from 15 videos) and we added two points on each side of the mouse where the hind legs connect to the torso. Tracking results were stored in CSVs.

Computational analysis of the tracking data as generated above were analyzed using the Keypoint MoSeq pipeline available here: https://dattalab.github.io/moseq2-website/index.html with adjustments to run with a GPU on Collab Enterprise of Google Cloud. Extraction was performed using the default configuration parameters and a select kappa value (kappa = 30,000,000). After generating a robust model as outlined in the pipeline, 44 syllables were produced for the *Grin2a* model, and 60 syllables were generated for the *Akap11* model. The number of frequencies of usage and number of transitions between syllables were then quantified. For visualization purposes, the usage plots only include syllables with a frequency above 0.5%. These syllables were manually annotated into five larger groups of behavior (grooming, pausing, darting, rearing, turning) and frequency of each behavioral group was quantified.

### Microdialysis

Microdialysis experiments were completed by Charles River Laboratories. L-shaped microdialysis probes with a polyacrylonitrile membrane (NO-PAN 6/2) were stereotaxically implanted into mice under anesthesia. The specific coordinates used for the striatum were (AP +0.8 mm, ML −1.7 mm from bregma, DV −3.5mm). 17 mice were used for microdialysis experiments (including 9 WT and 8 *Grin2a^+/-^*).

On Day 1 on experiments, the probes were perfused with artificial cerebrospinal fluid (aCSF, containing 147 mM NaCl, 3.0 mM KCl, 1.2 mM CaCl2, and 1.2 mM MgCl2) at a flow rate of 1.5 uL/min. Samples were collected over 30-minute periods by an automated fraction collector into vials containing 15 uL of stabilizing solution (0.2M formic acid and 0.04% ascorbic acid). The protocol includes a 2-hour pre-stabilization period, followed by 2 hours of baseline sampling, acute dosing (time = 0) and 4 hours of follow-up sampling. Locomotor activity was accessed simultaneously using a photobeam detection system (San Diego Instruments) in the home cage. At the study’s conclusion, animals were euthanized for probe placement verification via gross histology.

Dialysate samples were stored at −80 °C until analysis by High-Performance Liquid Chromatography coupled with tandem Mass Spectrometry (HPLC – MS/MS). The analysis utilized stable isotope (D_4_-DA) as an internal standard and involved SymDAQ derivatization. Analytes were separated on a reversed phase column (Phenomenex Synergi MAX-RP) and detected by an API-4500 triple quadrupole MS in positive ionization mode (Applied Biosystems, USA). Sample concentrations for analytes were determined using weighted (1/*χ*) regression against suitable in-run calibration curves using the Analyst data system (Applied Biosystems).

### Viruses

GRAB-DA and GCaMP sensors were purchased from Addgene, prepared as AAV9 viruses (no. 140554, 100845, 107790). The final virus constructs and concentrations used for this study were AAV9-hsyn-GRAB_DA2h (2.50 *x* 10^13^*gc*/*ml*), AAV9.Syn.Flex.GCaMP6s.WPRE.SV40 (2.10 *x* 10^13^*gc*/*ml*), and AAV9.CaMKII.GCaMP6s.WPRE.SV40 (1.90 *x* 10^13^*gc*/*ml*).

### Stereotaxic Surgery and Virus Injection

Mice were anesthetized with isoflurane and underwent injection of a 500 nanoliter injection of the selected virus (GRAB-DA, Flex-GCaMP, or CaMKII-GCaMP). To maintain consistency across animals, all injections were given on the left side. Virus injection was conducted using coordinates for the Dorsomedial Striatum (AP +0.50, ML +1,7 from bregma, DV −2.5 from brain surface). Optical fibers (MFC_400/430-0.66_3.0mm_MF2.5_FL from Doric Lenses) were placed on the left side ∼1mm above the injection site.

### Fiber Photometry

One hundred-two 12–20-week-old *Grin2a, Akap11 and wild-type* animals of mixed genotypes were used for stereotaxic surgeries and subsequent fiber photometry experiments. Only male mice were used for experiments. Following virus injection and fiber implantation, there was a three-week break to allow for full virus expression and surgical healing before starting recordings. All fiber photometry was recorded at 10kHz using the TDT fiber photometry system to record from both the 465 (GRAB-DA or GCaMP) and 405 (Isobestic) channels. Animas were renumbered for data collection and analysis to blind the experimenter to the genotype of the mice.

Before experiments, animals were habituated to the recording room and fiber photometry cables for a minimum of three days. Following adequate habituation, animals underwent a thirty-minute recording in their home cage to measure baseline activity (data not shown). During subsequent measurements, animals were allowed to move freely in an open field (40×40 cm) for recordings ranging from 15-60 minutes.

#### Signal Correction

Custom MATLAB scripts were developed for correction and normalization of the fiber photometry signal. Both the signal of interest and isosbestic recordings were down sampled to 10.17 Hz using local averaging. To correct for photobleaching of the signal, we fit a double exponential model (exp2) to both streams of signal. The raw signal was divided by then divided by this exponential fit. For motion correction, a linear regression (polyfit) was performed, regressing the bleach-corrected 465 signal to the isosbestic signal. The fitted signal was then subtracted from the bleach-corrected 465 channel. The final change in fluorescence was calculated by the following formula:

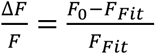

#### Comparison of Event Characteristics

For event-identification, a rolling z-score was applied to the dF/F signal using a causal sliding window of 20s. To prevent large events from artificially inflating the mean, the local mean was estimated using the 10^th^ percentile of the window data. For all event-related analysis, the findpeaks function in MATLAB was used to identify calcium or dopamine events with a prominence of Z-score > 2 and a minimum distance between events of 5s.

To align events with different velocity bins. Top-down video recordings were synchronized with fiber photometry recordings. To track velocity, a DeepLabCut model was used. We calculated percentile thresholds based on all the animals’ velocities in the experiment and divided these into equally sized bins, with the lowest velocity observed being 0 cm/sec and the highest being 35 cm/sec. Using the findpeaks function and a prominence threshold (Z > 2), we identified events in each of these bins for each animal.

In analyses aligning velocity to dopamine or calcium events, we used a similar event detection (findpeaks) to identify events with a specific prominence (Z > 2) and a minimum distance between of 5s. For each detected peak, the corresponding velocity trace was extracted in a window around the event of [−1, 2s]. For each animal, the velocity traces associated with the events were averaged and normalized to the 1s window immediately preceding the event onset.

#### Alignment of signal with movement onset and offset

To capture movement onsets, the velocity signal was first smoothed using a moving mean and a fixed time window of 1.67 seconds. Then, a changepoint detection algorithm was applied to the velocity derivative and onset indices were determined by identifying peaks (findpeaks function) in the changepoint-score (square of the velocity derivative, acceleration proxy). The minimum prominence of a peak was one standard deviation away from the mean of changepoint-scores. An onset was accepted only if the maximum velocity reached within the subsequent 1.5 seconds exceeded a dynamic threshold of the 80^th^ percentile of all velocity data collected across animals in the experiment. Accepted onsets were also required to be at least 2 seconds apart.

To measure movement offsets (pauses), a distinct detection method was implemented. First, a dynamic velocity threshold was established by calculating the 50^th^ percentile of the aggregated velocity dataset. Potential movement offset points (changepoints) were identified as the start index of any consecutive block of frames where the smoothed velocity fell below this threshold and lasted for a minimum of 0.5s. Accepted movement offsets had to be at least 2 seconds apart. To compare dF/F signals at movement onsets and offsets across animals, the analysis used a bootstrapping approach. dF/F signal was used rather than z-score to allow for capture of absolute signal amplitude. For each animal, the full valid set of traces aligned to the movement onset or offset was used to generate 1000 bootstrap iterations. In each iteration, a fixed number of traces (capped by the animal with the fewest valid traces) were randomly sampled with replacement and a mean trace was calculated. The final subject average was taken as the mean of all 1000 bootstrapped traces. The area under the curve was calculated (using the trapezoid function) directly from this stable, bootstrapped mean.

#### Light and Tone Presentations

For the light and tone presentations, fiber photometry recordings were conducted in Med Associate’s Operant Chambers (8.5” x 7.12” x 5”). The chambers had holes drilled through their tops to allow for fiber photometry cables to connect to animals while enclosed in the chamber. On the day of testing, mice were brought into the testing room thirty minutes prior to experiments and allowed to habituate to the boxes for five minutes prior to testing. Light (55 Lux, 200 ms) were given 20 times and tone (60 dB, 2 sec) presentations were given 20 times with 1 minute in between trials. For analysis of averaged dopamine response (dF/F) during the cue presentation, the signal was aligned to the time of the cue onset and normalized to [−2 to −1s] prior to the T = 0. The area under the curve was calculated (using the trapezoid function) for each trial and averaged by subject for each block.

#### Object Interaction

Object interaction was measured in fiber photometry animals with the same protocol and behavioral analysis as described above. Top-down video recordings were synchronized with fiber photometry recordings to allow for quantification of signal related to approach-related behavior. For each approach, the dF/F signal was aligned to the time the mouse entered the object zone (2 cm from object perimeter) and normalized to the average window of [−2 to −1s] prior to zone entry. The area under the curve was calculated (using the trapezoid function) for each approach and averaged by subject.

### Virus Injection Site Validation

Following experiments, mice were euthanized under isoflurane with transcardial perfusion of ∼25 ml of PBS followed by ∼25 ml of 4% paraformaldehyde (PFA). Coronal brain sectioning of slices near the optical fiber implants was performed using a vibratome (Leica VT200s). The brain sections were mounted onto slides using Vectashield Antifade Mounting Media with DAPI (Vector Laboratories, H-1200-10) and fluorescence of GRAB-DA and GCaMP sensors was imaged with a Zeiss Axioscan 7 slidescanner.

### Statistics

Statistics for behavior and fiber photometry experiments were calculated using GraphPad Prism (version 10), with each marker representing the mean ± SEM of an individual animal. For comparisons between multiple groups (e.g. across genotypes or velocity bins), a two-way or one-way Analysis of Variance (ANOVA) was used, followed by an appropriate post-hoc test (Tukey’s multiple comparisons test). When comparing only two independent groups, an unpaired t-test was applied; if the assumption of equal variances was violated, a Welch’s t-test was used. For data sets where assumptions for parametric testing (normality or homoscedasticity) were not met, the appropriate nonparametric equivalent (e.g. Kruskal-Wallis test for multiple groups) was employed. The specific test used for each comparison is indicated in the figure legends. Significance was set at p < 0.05 except for Keypoint MoSeq syllable usage plots which used a threshold of FDR < 0.1 for significance. The histological validation of viral expression and optical fiber placement in the DMS is shown in **Extended Fig 8**.

## Supplementary Figures

**Extended Figure 1:**
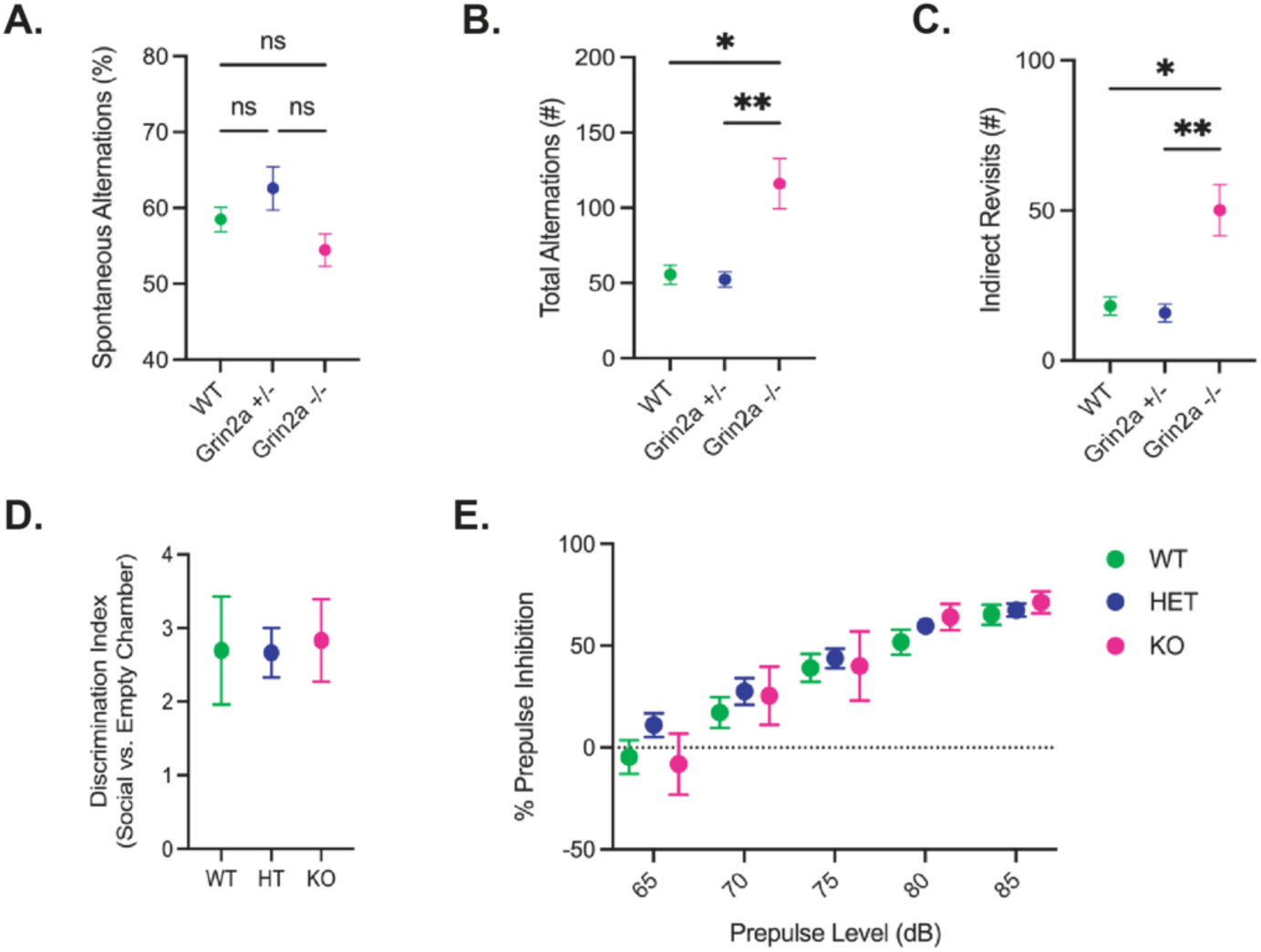
**A.** Percentage of correct spontaneous alternations in Y-maze test. **B.** Total number of alternations in Y-maze test. **C.** Total number of indirect revisits in the Y-maze test. **D.** Ratio of time spent in chamber with mouse (social) vs. time spent in chamber without mouse (empty) in a three-chamber social interaction test. **B,C,D,.** Kruskal-Wallis test with Dunn’s multiple comparisons test.. *, p < 0.05; **, p < 0.01. n = 6 WT, 7 *Grin2a^+/^*^-^, 8 *Grin2a^-/-^* mice. **E.** Percentage of prepulse inhibition at different decibel levels. No significant differences between genotypes (Repeated measures two-wave ANOVA with Tukey’s post hoc test). n = 10 WT, 17 Grin2a^+/-^, 9 Grin2a^-/-^.

**Extended Figure 2:**
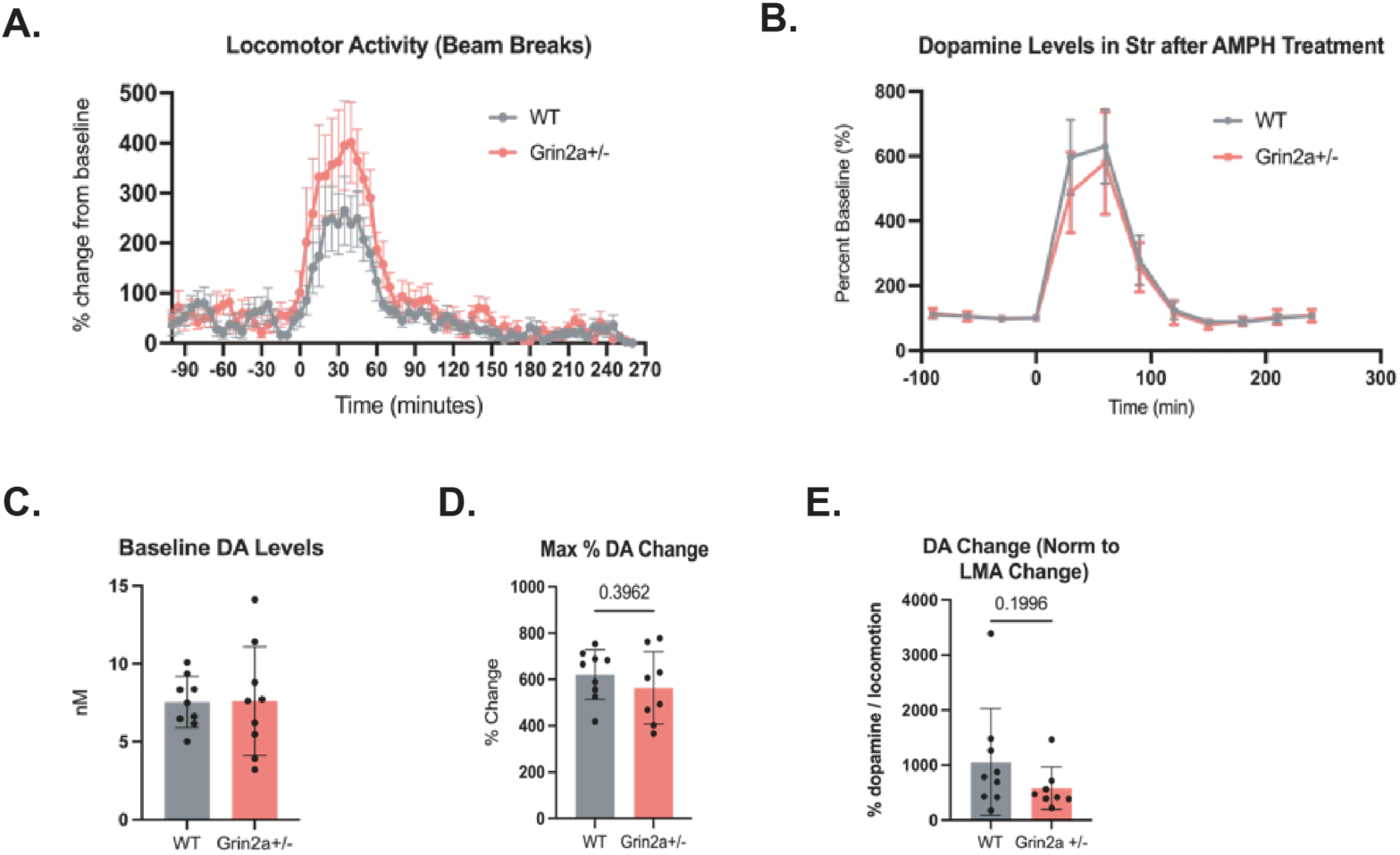
**A.** Average change in locomotor activity (LMA) from baseline during AMPH treatment. Using two-way ANOVA, a significant effect of AMPH treatment was observed (p<0.0001) and of the interaction of AMPH treatment and genotype (p = 0.0355), but no significant effect of genotype alone was found (p =0.449). Tukey’s post hoc analysis did not reveal significant genotype effects at any specific timepoints. N = 9 WT, 8 *Grin2a^+/-^*. **B.** Average change in dopamine levels from baseline (T=0), measured by microdialysis, during amphetamine (AMPH) treatment. Using a two-way ANOVA, a significant effect of AMPH treatment was observed (p<0.0001) but no effect of genotype was found (p = 0.0891). N = 9 WT, 9 *Grin2a^+/-^*. **C.** Maximum percent change in dopamine during AMPH treatment. No significant difference found (Welch’s t test, p = 0.3962). **D.** Maximum percent change in dopamine during AMPH normalized to the change in locomotor activity from pre-AMPH treatment (T = −90 to T=0) to post-AMPH treatment (T= 0 to T = 90).

**Extended Figure 3:**
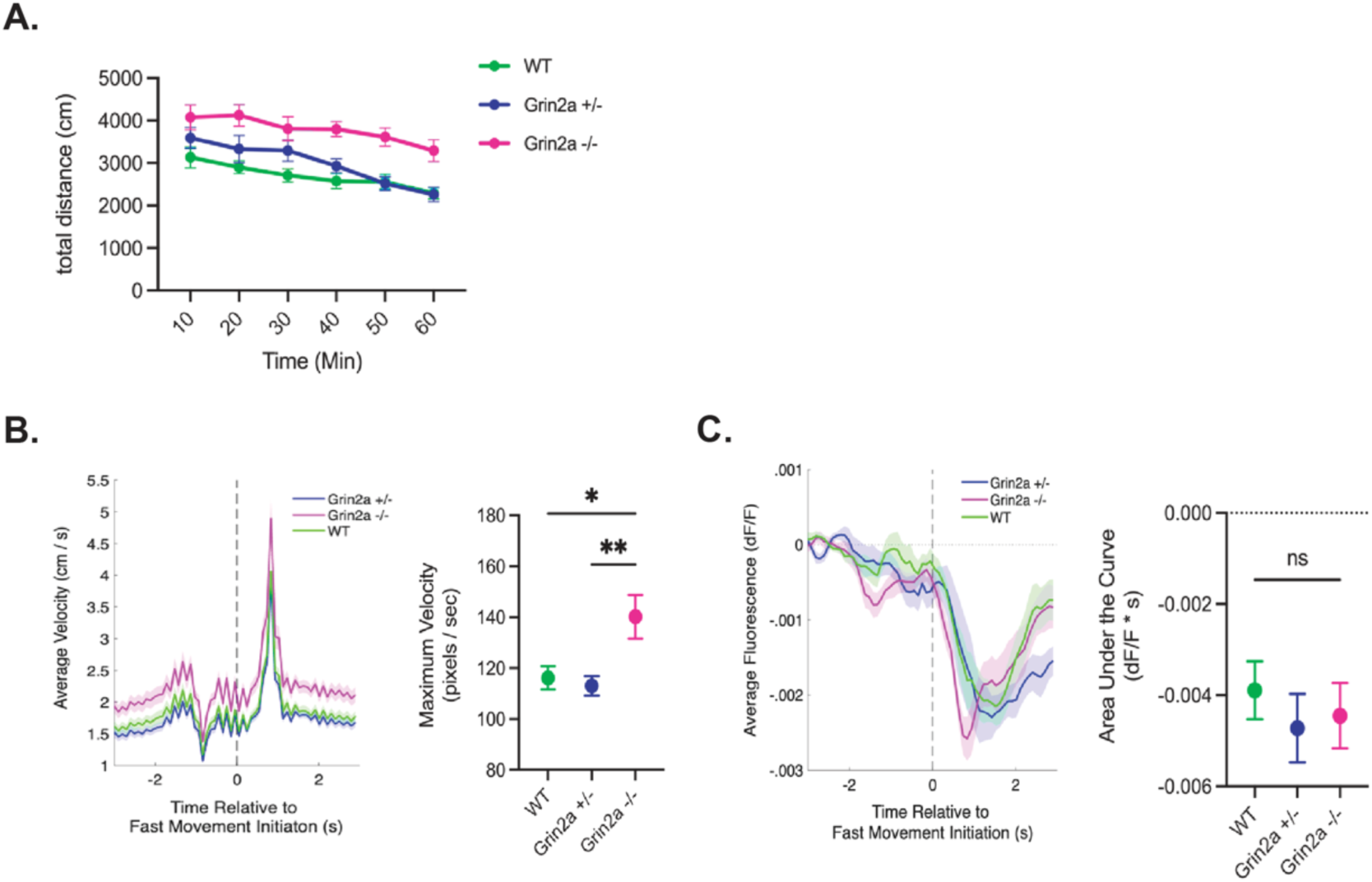
**A.** Distance traveled between WT, and *Grin2a* mutant, animals injected with GRAB-DA2h during 1-hour open field recording. N = 10 WT, 11 *Grin2a^+/-^*,11 *Grin2a^-/-^*. **B.** Average velocity of WT and *Grin2a* mutants during a movement onset. Maximum achieved velocity was quantified between genotypes (One way ANOVA, Tukey’s multiple comparison test). *, p < 0.05; **, p < 0.01. **C.** Average dopamine signal of WT and *Grin2a* mutants during a movement onset. Area under the curve was calculated from 0-3s following movement onset.

**Extended Figure 4:**
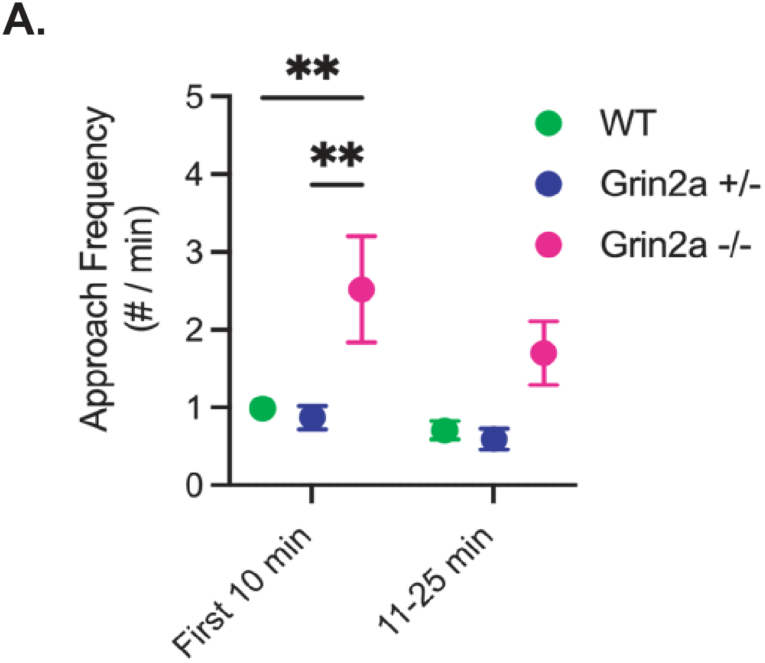
**A.** Approach frequency during object interaction recordings in GRAB-DA injected WT and *Grin2a* animals. Frequency was divided between the first 10 minutes of the test and 11-25 minutes. To compare genotypes at both timepoints, a repeated measures 2-way ANOVA with Tukey’s post hoc test was performed. *, p < 0.05; **, p < 0.01.

**Extended Figure 5:**
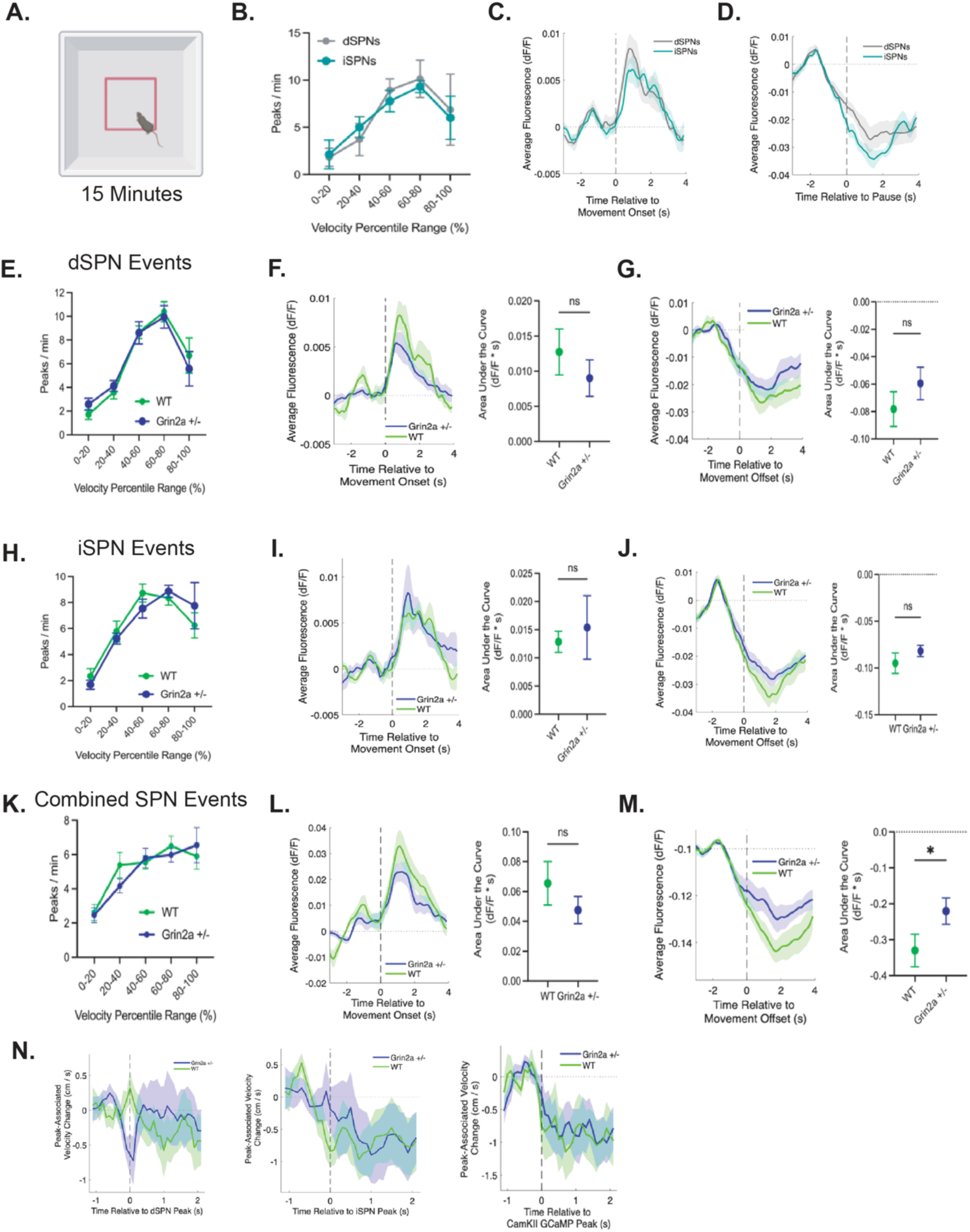
**A.** Schematic of open field recording. **B.** Comparison of peak frequencies for dSPNs and iSPNs over different velocity percentile bins (N = 6 Drd1-Cre and 7 A2A-Cre). **C.** Average waveforms of dSPNs and iSPNs plotted during movement onset. **D.** Average waveforms of dSPNs and iSPNs plotted during movement offset. **E.** Comparison of dSPN peak frequencies between WT and *Grin2a +/-* over different velocity percentile bins. N = 6 WT and 7 *Grin2a^+/-^*. **F.** Average dSPN signal in WT and *Grin2a^+/-^*animals during movement onset. Comparison of area under the curve (calculated from 0 to 4s post movement onset) between genotypes. **G.** Average dSPN signal in WT and *Grin2a^+/-^*animals during movement offset. Comparison of area under the curve (calculated from 0 to 4s post movement offset) between genotypes. **H.** Comparison of iSPN peak frequencies between WT and *Grin2a +/-* over different velocity percentile bins. N = 7 WT and 7 *Grin2a^+/-^*. **I.** Average iSPN signal in WT and *Grin2a^+/-^*animals during movement onset. Comparison of area under the curve (calculated from 0 to 4s post movement onset) between genotypes. **J.** Average iSPN signal in WT and *Grin2a^+/-^*animals during movement offset. Comparison of area under the curve (calculated from 0 to 4s post movement offset) between genotypes. **K.** Comparison of total DMS bulk calcium peak frequencies from all SPNs between WT and *Grin2a +/-* over different velocity percentile bins. N = 9 WT and 10 *Grin2a^+/-^*. **L.** Average bulk calcium signal from all SPNs in WT and *Grin2a^+/-^* animals during movement onset. Comparison of area under the curve (calculated from 0 to 4s post movement onset) between genotypes. **M.** Average bulk calcium signal from all SPNs in WT and *Grin2a^+/-^* animals during movement offset. Comparison of area under the curve (calculated from 0 to 4s post movement offset) between genotypes. **N.** Event (peak) associated velocity changes for iSPN, dSPN, and combined SPN calcium recordings in WT and *Grin2a+/-* animals

**Extended Figure 6:**
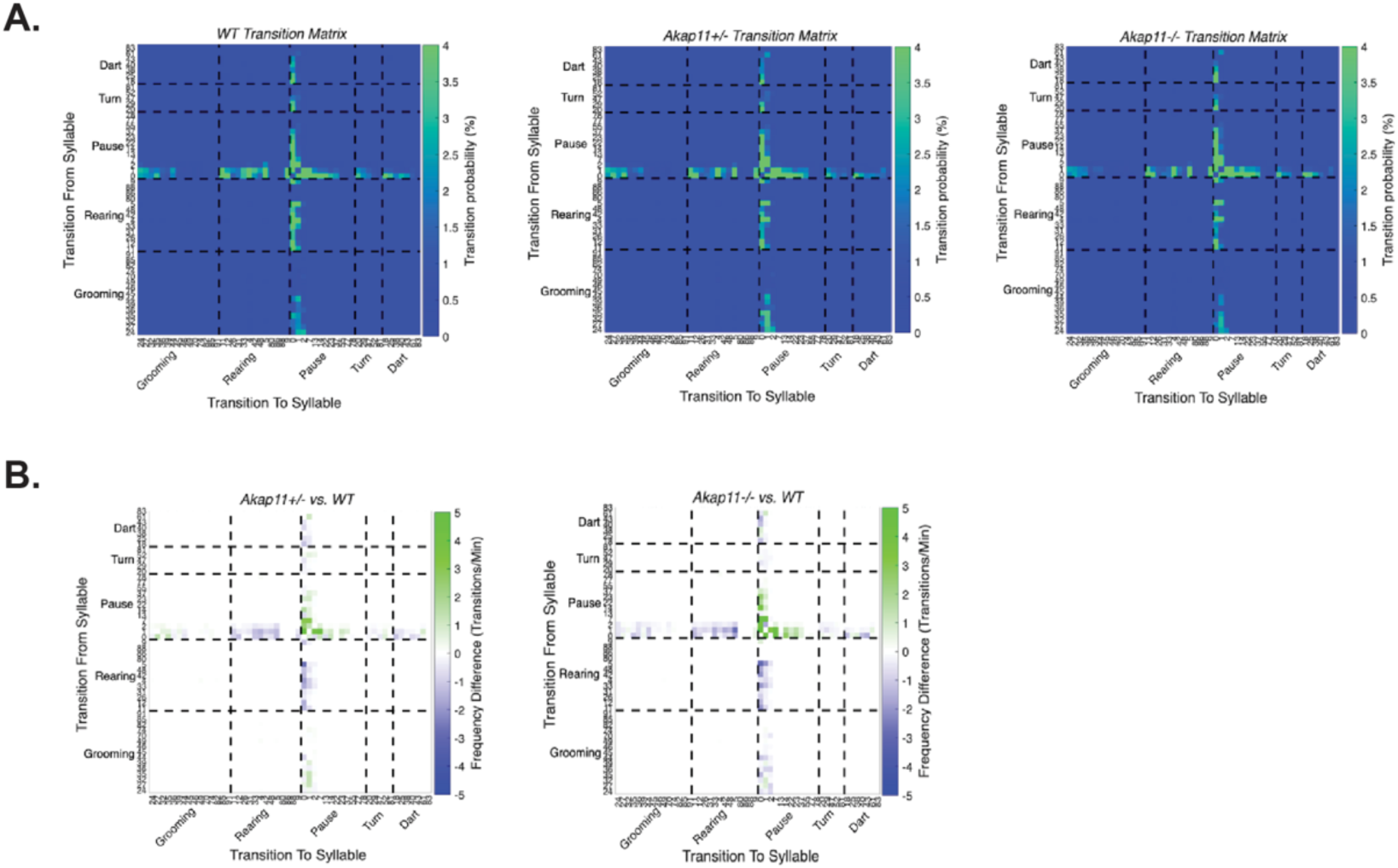
**A.** Heat map of transition matrices between syllables for WT, *Akap11^+/-^*, and *Akap11^-/-^* animals, color bar represents scale of probability of transition. Syllables re-ordered according to behavioral groups. **B.** Heat map of subtraction of transition probabilities between WT vs. *Akap11^+/-^* and WT vs. *Akap11^-/-^*.

**Extended Figure 7:**
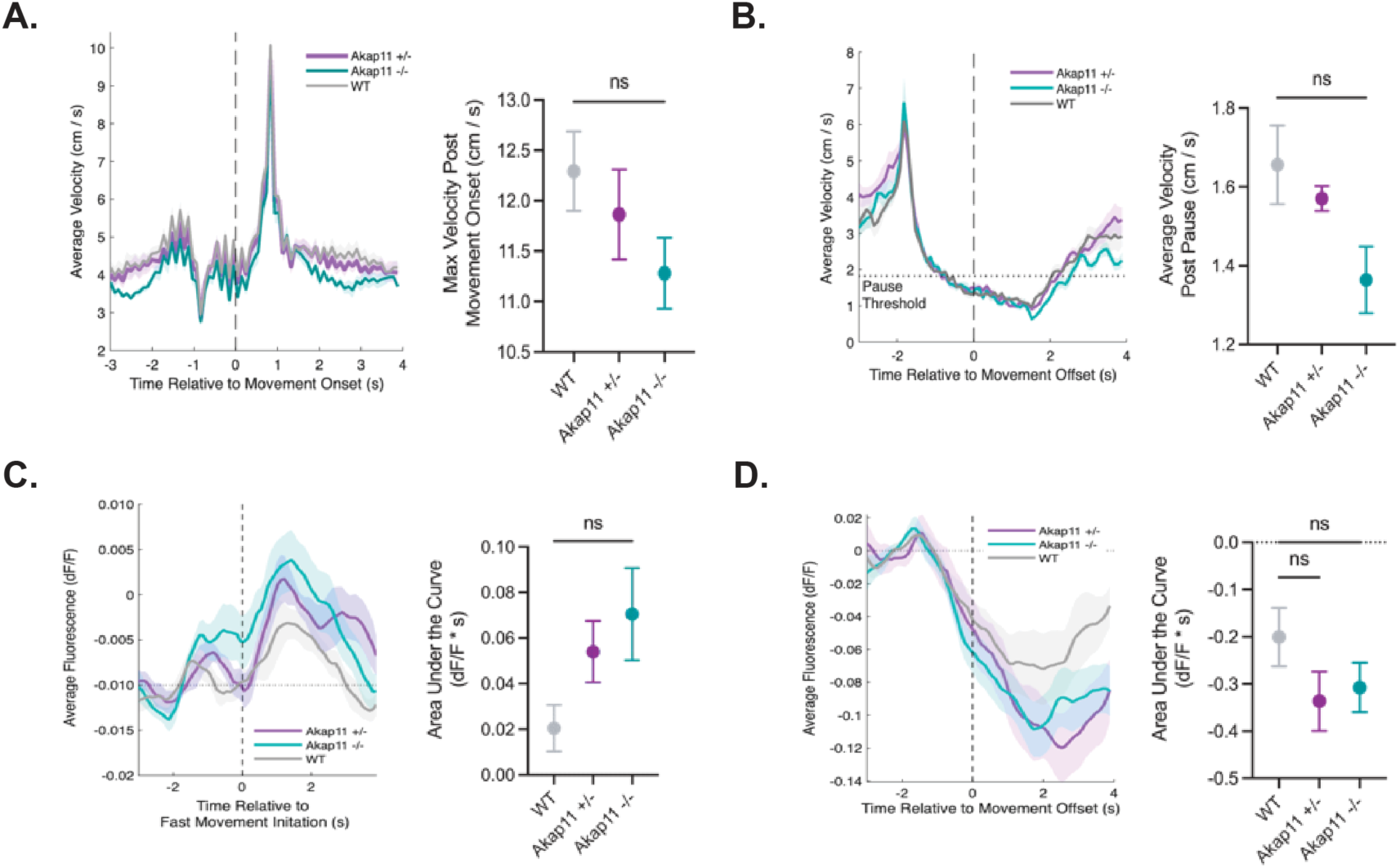
**A.** Average velocity of WT and *Akap11* mutants during a movement onset. Maximum achieved velocity was quantified between genotypes. **B.** Averaged velocity for WT and *Akap11* mutants over time during movement offset. Average velocity post-pause is shown with SEM for each genotype. **C.** Average dopamine signal of WT and *Akap11* mutants during a movement onset. Area under the curve was calculated from 0-3s following movement onset. **D.** Average dopamine signal of WT and *Akap11* mutants during a movement offset. Area under the curve was calculated from 0-3s following movement offset. **A, B, C,D.** One way ANOVA, Tukey’s multiple comparison test.

**Extended Figure 8:**
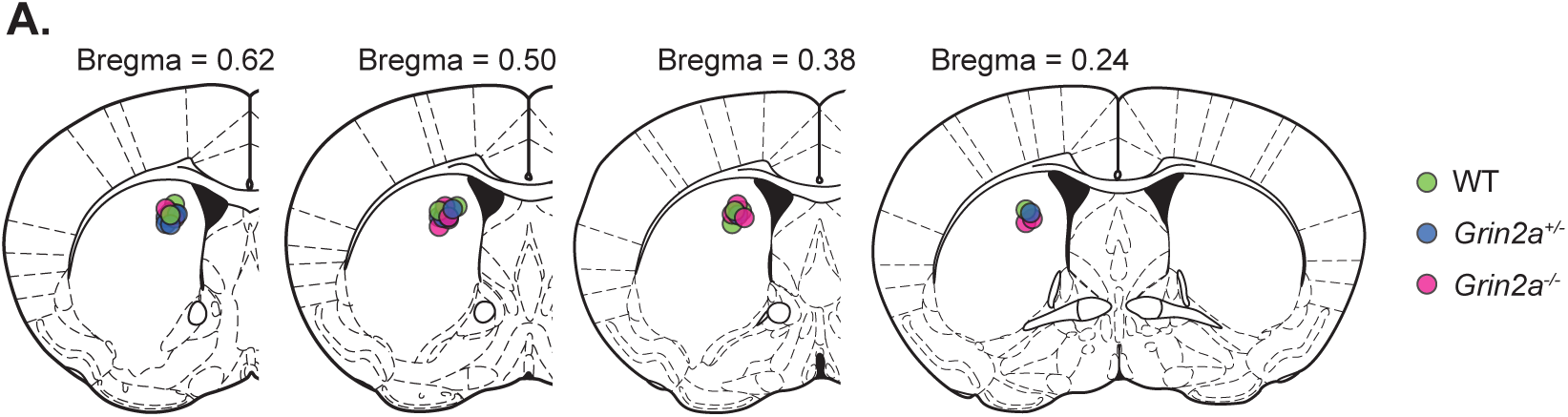
A. Schematic coronal brain sections showing the optical fiber tip locations in GRAB-DA injected WT (n=10), *Grin2a^+/-^* (n=11), and *Grin2a^-/-^* (n=11) mice used in the experiments described in Figure 3. Each dot represents an individual optical fiber placement. The number above each section represents A-P coordinate relative to bregma (mm).

